# The Enderstruder: An accessible open-source syringe extruder compatible with Ender series 3D printers

**DOI:** 10.1101/2023.08.15.553241

**Authors:** Domenic J. Cordova, Angel A. Rodriguez, Sabrina C. Woodward, Cody O. Crosby

**Author notes:** **Corresponding author’s email address and Twitter handle**, @crosby_sw.

## Abstract

Bioprinting has enabled the precise spatiotemporal deposition of cells and biomaterials, opening new avenues of research in tissue engineering and regenerative medicine. Although several open-source syringe extruder adaptations for bioprinters have been published and adopted by end users, only one has been specifically adapted for the Ender series, an affordable and open-source line of thermoplastic 3D printers. Here, we introduce the Enderstruder, a cost-effective extruder attachment that uses a standard 10 mL BD syringe, positions the stepper motor at the level of the gantry, enhances x-axis stability with a linear rail, and uses the originally included stepper motor, resulting in reduced cost and simplified assembly. Furthermore, we present an iterative process to fine-tune printing profiles for high-viscosity biomaterial inks. To facilitate the implementation of our work by other researchers, we provide fully editable Cura profiles for five commonly used biomaterials. Using these five materials to validate and characterize our design, we employ the Enderstruder to print established calibration patterns and complex shapes. By presenting the Enderstruder and its iterative development process, this study contributes to the growing repository of open-source bioprinting solutions, fostering greater accessibility and affordability for researchers in tissue engineering.

**Specifications table:** 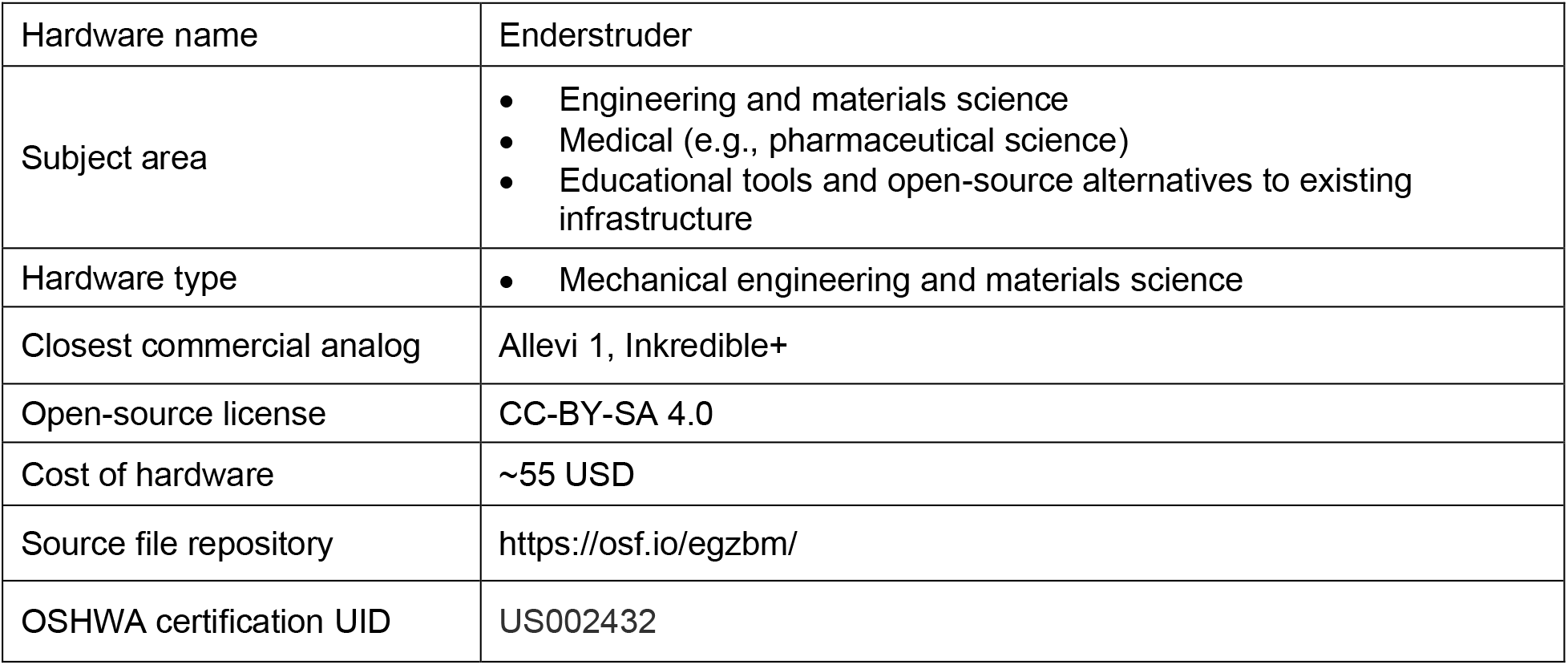

## 1. Hardware in context

The 3D printing industry has experienced remarkable growth in the past decade. This growth and the expiration of key patents have led to the democratization of this advanced additive manufacturing technology, resulting in a flourishing market for open-source, low-cost 3D printers that have been shown to have myriad applications in the modern laboratory [1]. Open-source machines and software have enabled the fabrication of traditional thermoplastics (e.g., polylactic acid (PLA)); however, traditional thermoplastics are not suitable mimics of the human body’s extracellular matrix (ECM) and are therefore not suitable for cytocompatible tissue scaffold manufacturing. Bioprinting, an additive manufacturing technology defined by the spatiotemporal deposition of soft biocompatible polymers exhibiting near-fluid properties at room temperature, presents unique challenges for conventional fused filament fabrication (FFF) 3D printer technology [2]. Although commercial bioprinters employing extrusion, vat polymerization, and droplet-based techniques have made it to market [3], they are expensive and proprietary, with high price tags and closed-access software. As a result, more laboratories may increasingly rely on open-source, inexpensive, and highly customizable bioprinter designs [4]. In this study, we specifically focused on an extrusion-based approach because extruder 3D bioprinter adaptations are affordable, easily constructed, and accessible to scientists with minimal hardware or software design experience.

Extrusion-based 3D bioprinters can be developed by scratch-building the motion and extrusion system or designing novel syringe extruders for biological applications. Scratch-built bioprinters showcase impressive engineering [5–8] but often fail to surpass the performance of cost-effective 3D positioning systems available in the market, such as the Anet A8 and Ender series printers. However, these scratch- built solutions could be viable alternatives for resource-constrained labs and valuable learning experiences for students seeking hardware and software fundamentals expertise. Designing a syringe extruder that can be mounted to an existing 3D motion system is often simpler, optimizing affordability and performance. A landmark paper by Wijnen *et al.* led to the development of the first reliable open-source syringe pump system [9]; two other publications rapidly adapted this syringe pump for bioprinting applications [10,11]. However, these designs are either "top-heavy" or introduce considerable dead volume into the system. In a previous review, we surveyed the literature and found eight open-source syringe extruders (five published in *HardwareX*) that were replicable. We compare these eight designs in chronological order in **Table 1** to the syringe extruder described in this manuscript, the Enderstruder.

**Table 1:** "Cost" excludes the 3D printer and the PLA/ABS filament, "Dead volume" refers to tubing or connectors that lead to loss of material, "Adds torque" means that the stepper motor’s torque is supplemented by a gearbox or geared system, "Top-heavy" indicates that the stepper motor, which makes up the bulk of the weight, is positioned significantly above the gantry.

| Authors (Design Name) [Citation] | Hardware X | Cost (USD) | Dead volume | Retraction-enabled | Original stepper motor | Adds torque | Syringe volume | Top-heavy |
| --- | --- | --- | --- | --- | --- | --- | --- | --- |
| Pusch *et al.* (Large-Volume Extruder, LVE) [12] | Yes | 49 | Yes | Yes | Yes | Yes | 60 mL BD | No |
| Bessler *et al.* (Nydus One Syringe Extruder, NOSE) [13] | Yes | 90 | No | No | Yes | No | 2/5/10 mL BD | Yes |
| Kahl *et al.* [14] | No | 159 | No | Yes | Yes | No | 1 mL BD | No |
| Tashman *et al.* (Replistruder) [15] | Yes | 62.9 (with 5 mL Hamilton) | No | Yes | Yes | Yes | 10 mL BD, 2.5/5 mL Hamilton | Yes, without mount re-design |
| Darling and Smith (SPECS) [16] | Yes | 277 | Yes | Yes | Yes | No | 5/60 mL BD | No |
| Dávila *et al.* (iCAN-X) [17] | No | 91 | No | Yes | Yes | No | 10 mL BD | No |
| Chimene *et al.* (MOSB) [18] | No | 400 | No | Yes | No | Yes (with gearbox) | Makin clay gun extruder | Yes |
| Demircan and Özçelik [19] | Yes | 22 | No | Yes | Yes | No | 60 mL BD | No |
| Cordova *et al.* (Enderstruder) | Yes | 55 | No | Yes | Yes | Yes | 10 mL BD | No |

Most designs listed in **Table 1** either leave significant material in the connective tubing, do not add additional torque to the usually underpowered extruder motors, or position the stepper motor significantly above the gantry, introducing positioning errors at high printing speeds. Of these designs, the Replistruder, produced by Prof. Adam Feinberg’s group at Carnegie Mellon University, remains the most widely adopted open-source design within the bioprinting community, with 15 manuscripts incorporating the Replistruder in their Methods section. Compared to the Replistruder, the Enderstruder

- Has a lower center of gravity. The Replistruder’s center of gravity could be lowered, though this would require re-designing the mount.
- Exhibits customizable 3D-printed gears at a 4:1 ratio (offering 33% more torque than the Replistruder’s 3:1 gearing)
- Does not require belt tensioning as the 3D-printed gears are rigid
- Enables a more facile exchange of the syringe
- Supports the syringe and limits syringe barrel movement

Notably, most open-source designs are optimized for Prusa machines with a belt-driven mechanism mounted on two smooth cylindrical rods. We compared 28 open-source extrusion 3D bioprinters and found six explicitly designed for Prusa 3D printers. The only thermoplastic printers to be included in multiple manuscripts were Anet A8 (3), MakerBot Replicator 2X (2), and Printrbot Simple Metal (2), which is no longer manufactured. As the naming convention implies, the Enderstruder is designed explicitly for Ender series machines. Among the popular open-source 3D printers available in 2023, the Ender (Creality^TM^) family, including the Ender 3, Ender-3 Pro, and Ender 3 V2, stand out as affordable open-source options with many accessible video tutorials on YouTube [20]. These printers, often found for less than 200 USD, use a length of aluminum v-slot extrusion as the primary x-axis gantry, with three rubber wheels to allow for movement along the extrusion. In *HardwareX* alone, Ender-series printers have been modified for diverse uses in the biological sciences, including histological staining [21] and a programmable syringe pump set [22].

To our knowledge, only two open-source syringe extruders, developed by Sun and To [23] and Chimene *et al.* [18], have been designed to be easily incorporated into Ender machines. The former mounts an open-source paste extruder, the "Spritzstruder," to the frame and connects the needle to the extruder via a long piece of flexible tubing, which results in considerable dead volume. The syringe extruder published by Chimene *et al.*, designed as a modified clay gun extruder, is mounted on a vertical linear rail, and incorporates temperature control and automated bed leveling features. The design is robust and can print complex geometries, but substantial firmware changes must be made to operate the syringe extruder. The clay gun extruder has a large diameter and is not well-suited for small volumes; it is also opaque, and it is not easy to gauge whether the remaining material remains in a sufficient and printable state. Gusmão *et al.* recently published an Ender 3 V2 modification for multiple extruders equipped with temperature control and UV cross-linking; however, the design is not open-source, and it is difficult to evaluate the efficacy of the novel syringe extruder [24]. Our objective was to create an open-source extruder for Ender 3D printers that could be utilized by all researchers interested in the bioprinting discipline, even those with limited hardware and software design experience. With a focus on simplicity, we have introduced a design that uses a standard 10 mL BD syringe instead of a more expensive clay gun extruder, lowers the center of gravity of the system by mounting the motor at the level of the gantry, adds x-axis stability with a linear rail, and uses the originally included stepper motor, driving down the price and ease of assembly.

## 2. Hardware description

Our syringe extruder consists of four primary components: a 3D-printed core, a 3D-printed syringe carriage, a stock motor, and a readily available, inexpensive syringe (**Figure 1**). The core connects the mounted extruder to the Ender’s x-axis gantry, ensures system rigidity, and prevents needle deflection when extruding highly viscous biomaterial inks. It is attached to the rail carriage and motor using M3 screws, incorporates slots for cylindrical linear rails, and features slits for the toothed belt, facilitating translation along the x-gantry. The NEMA 17 stepper motor, included with most Ender machines, drives a threaded rod connected to a 3D-printed gear system with a 4:1 ratio. The syringe carriage links the Carriage and the Core, transmitting the geared system’s torque.

**Figure 1:**
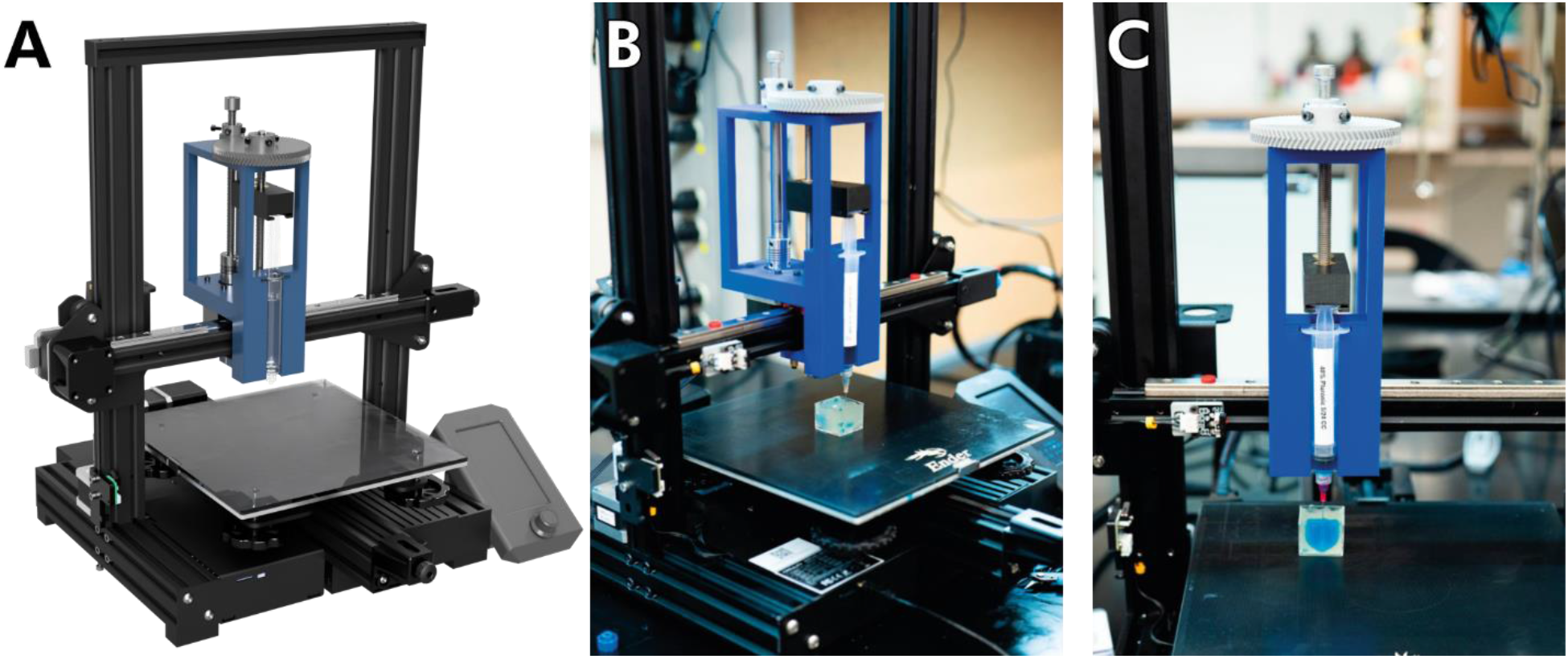
CAD models and photographs of the Enderstruder **(A)** A 3D render completed in Autodesk Fusion 360 of an Enderstruder mounted to an Ender 3 V2 and **(B)** a left-side and **(C)** front-side photograph of the Enderstruder.

Unlike many FFF 3D printers currently on the market, the Ender series features a rolling carriage mounted on a bar of v-slot extrusion. To minimize dead volume introduced by additional tubing [16], we directly mounted the syringe extruder onto the gantry. We replaced the rubber wheels with a linear rail to **(1)** achieve smoother movement, **(2)** reduce ringing artifacts, and **(3)** stabilize the increased weight from the stepper motor. As noted by Demircan and Ozcelik, an unbalanced center of gravity around the linear rail can result in uneven layers and compromised 3D resolution [19]. Therefore, we positioned the motor at the gantry level, as close to the linear rail as feasible within specified tolerances.

The key objective was to ensure accessibility for individuals with limited hardware and software design expertise. To achieve this, we adopted several strategies. First, we minimized the number of 3D- printed components in our design, utilizing only five parts that most printers can produce within a single printing session of less than a day. Second, we maximized the utilization of original Ender components, particularly the extruder stepper motor, eliminating the need for additional wiring or extensive modifications to the existing Marlin firmware. Third, our 3D-printed core was designed to use a 10 mL Becton-Dickinson (BD) syringe, an affordable plastic syringe costing approximately $0.29 per unit on Fisher Scientific, an approved BD supplier.

Moreover, the core can be easily modified in any computer-aided design (CAD) program to accommodate syringes of different sizes or alternative designs, such as the Hamilton glass autoclavable syringes commonly employed in bioprinting studies. Fourth, we conducted extensive printing experiments with various biomaterial inks and have shared the corresponding open-source Cura profiles, enabling researchers to replicate the results presented in this paper. Currently, the creation of slicing profiles often relies on a "guess-and-test" approach or AI algorithms, which limit a comprehensive and mechanistic understanding of the impact of different printing parameters. This article details our iterative methodology for determining optimal flow rates for known and novel materials.

We believe our design and methodology will be of particular interest to:

● **Biomaterials scientists**. Those who design and test novel biomaterial inks need custom control of their machine and an in-depth understanding of the parameters affecting their novel biomaterial’s printability. Furthermore, our straightforward methodology for creating novel Cura profiles should reduce the time necessary for scaffold fabrication with a new material.
● **Laboratories with limited funds**. The printer and extruder have a combined cost of less than 260 USD and do not require component or software subscriptions; this allows the printer to be continually updated as hardware and software upgrades emerge on the 3D printing scene.
● **Laboratories without a background in the engineering disciplines**. We intentionally simplified the design to be accessible to those with no or limited mechanical and electrical engineering background.
● **Teaching laboratories at the undergraduate and graduate levels**. This design can be easily built at scale and be used to give students an in-depth experience with 3D printing, mechanical design, and polymer synthesis/characterization.

## 3. Design files summary

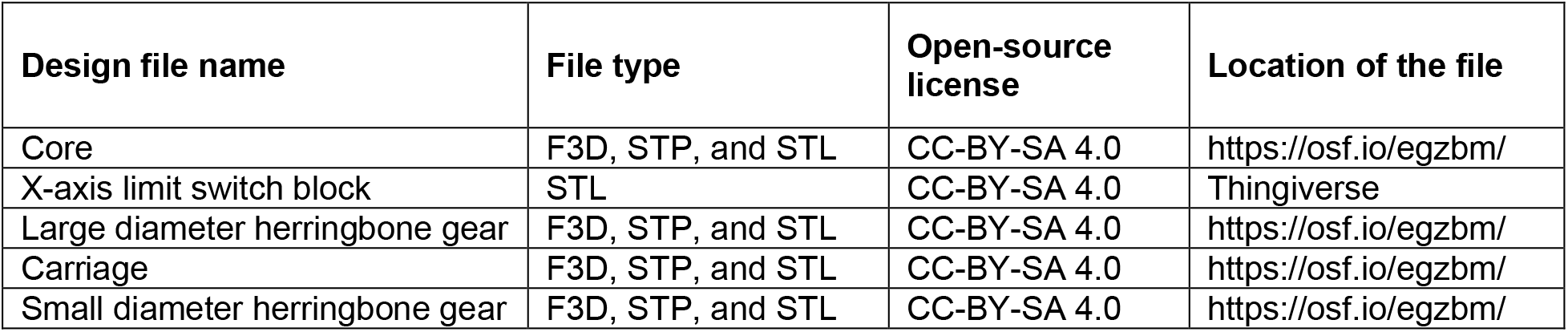

### Core

The core attaches the extruder to the linear rail and provides a rigid frame for the built-in syringe pump. The core should be printed with 30% gyroid infill and 0.2 mm layer height. We placed support material everywhere, used a 50% overhang angle, and printed support "lines" with a 5% or 10% density. The core was printed with the right or left side in contact with the bed, assuming the syringe holder represents the front orientation of the core. On our Ender series printer, we found that a retraction setting of 7.5 mm at 80 mm/s prevented noticeable stringing; this pair of settings can vary considerably among individual machines and should be calibrated beforehand. We have uploaded our generic PLA printing profile to the OSF repository for user convenience (**Figure 2A**).

**Figure 2:**
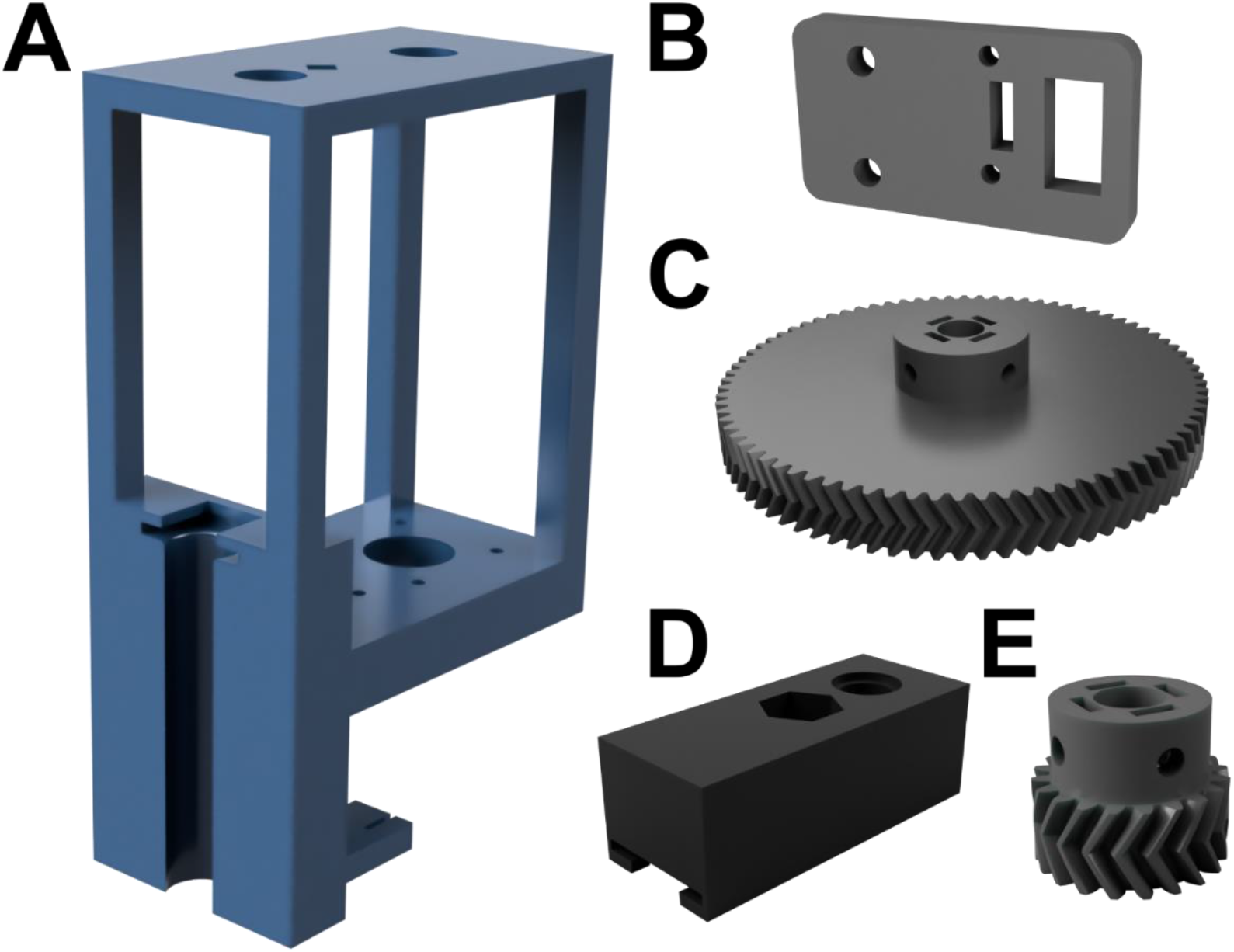
Renders of the 3D-printed PLA components **(A)** Core **(B)** Carriage **(C)** Small diameter herringbone gear **(D)** Large diameter herringbone gear. Please note that these renders are not to scale and are for illustration purposes only.

### X-axis limit switch block

This extends the x-axis limit switch to avoid a collision of the Enderstruder with the motor bracket. This block can be printed at 30% gyroid infill without support material (**Figure 2B**).

### Large diameter herringbone gear

This gear is attached to the carriage lead screw via four M3x6 socket-head screws (**Figure 2C**). Both gears were printed with the orientation shown in **Figure 2** at 100% infill for structural rigidity, and no support material was necessary for a high-resolution print.

### Carriage

The carriage translates rotation from the leadscrew to vertical movement along the z-axis. The carriage is rigid and has a slot for the top of the syringe handle. The Carriage was printed with the same settings as the core and the orientation shown in **Figure 2D**.

### Small diameter herringbone gear

This gear is attached to the motor leadscrew via four M3x6 socket- head screws. (**Figure 2E**)

## 4. Bill of materials summary

| Designator | Component | Number of units | Cost per unit (USD) | Total cost (USD) | Source of materials | Material |
| --- | --- | --- | --- | --- | --- | --- |
| M3 Screw and Nut Assortment<br>Hex Socket Head Cap | B0B51BFSWZ | 1 | 0.025 | 0.50 | Amazon | Metal |
| ½" Flanged Bearing | 6338K411 | 1 | 1.31 | 1.31 | McMaster-Carr | Metal |
| M8 Brass Nuts | 90690A055 | 2 | 0.90 | 1.80 | McMaster-Carr | Metal |
| Carriage Block and 300mm Linear Rail | MGN12H and MGN12 | 1 | 21.89 | 21.89 | Amazon | Metal |
| M3 Sliding T-Nuts | B08144SBGQ | 3 | 0.06 | 0.18 | Amazon | Metal |
| M8 x150mm Hex Socket Cap Threaded Rod | SSC-M8-150-A2 | 2 | 2.40 | 4.80 | ACUU | Metal |
| M8 Flanged Bearing | 6659K676 | 2 | 4.24 | 8.48 | McMaster-Carr | Metal |
| Flexible Shaft Coupling (M8-M5) | B01HBPHSII | 1 | 8.99 | 8.99 | Amazon | Metal |
| Linear Shaft ¼" Diameter, 5" Long | 6061K413 | 1 | 6.24 | 6.24 | McMaster - Carr | Metal |
| 1 kg of Plastic Filament | Grey PLA Hatchbox | ~300 g | 7.99 | 26.99 | Amazon | PLA |
| Thermoplastic 3D Printer | Ender 3 V2 | 1 | 199.00 | 199.00 | Creality | Metal |

## 5. Build instructions

### Removing and disconnecting the standard extruder head from an Ender series 3D printer

Please note that is demonstrated how to remove the extruder head from an Ender 3 V2 printer, though the process is nearly identical for other Ender series 3D printers.

WARNING: Before building the Enderstruder, please ensure that the printer is turned off and unplugged!

1. Remove the extruder head fan shroud by loosening two screws that fasten it to the translating carriage (**Figure 3A**). Once the fan shroud is removed, the locknut necessary to loosen the wheel will be accessible (marked in a red circle in **Figure 3B**).
2. Take the crescent wrench in the Ender series kit and secure the now accessible locknut in place while using the included hex key to loosen the metric screw until the wheel can be loosened (**Figure 3C**).
3. Disconnect the toothed belt by extracting it from the slots in the Carriage. The extruder head should now be able to be removed from the piece of horizontal extrusion. Now, the wiring needs to be disconnected from the mainboard.
4. A total of 5 screws must be removed on the bottom of the machine (3) (**Figure 4A**) as well as under the printer bed (2) (**Figure 4B, C**) to access the printed circuit board (PCB). Removing these screws will remove a panel and expose the necessary connections on the mainboard.
5. Remove the red/black wire connected to the extruder head cooling fan (**Figure 4D**). WARNING: Take a picture of the existing connection because it must be reconnected when you are finished with the following steps. You may notice hot glue on some of the connectors to keep them from loosening; this hot glue can be removed with the blue flush side cutters included with the Ender kit.
6. A total of 6 connections will need to be disconnected from the mainboard to remove the extruder head from the Ender series printer. The wires that will be disconnected come from the cable mesh; multiple zip ties may need to be cut to remove the cable mesh.
7. Locate the wire connections and disconnect each wire from the control board. Once the necessary wires have been removed, the extruder head can be removed from the machine.

a. The braided red, thin red and thin black connections must be unscrewed using a flathead screwdriver before they can be disconnected. (**Figure 4E**)
b. The red and blue connection and the two white wires on the far left of the control board can be pulled out. (**Figure 4F**)
8. As noted earlier, ensure that the cooling fan has been reconnected and re-fasten the five screws removed in Step 4.

**Figure 3:**
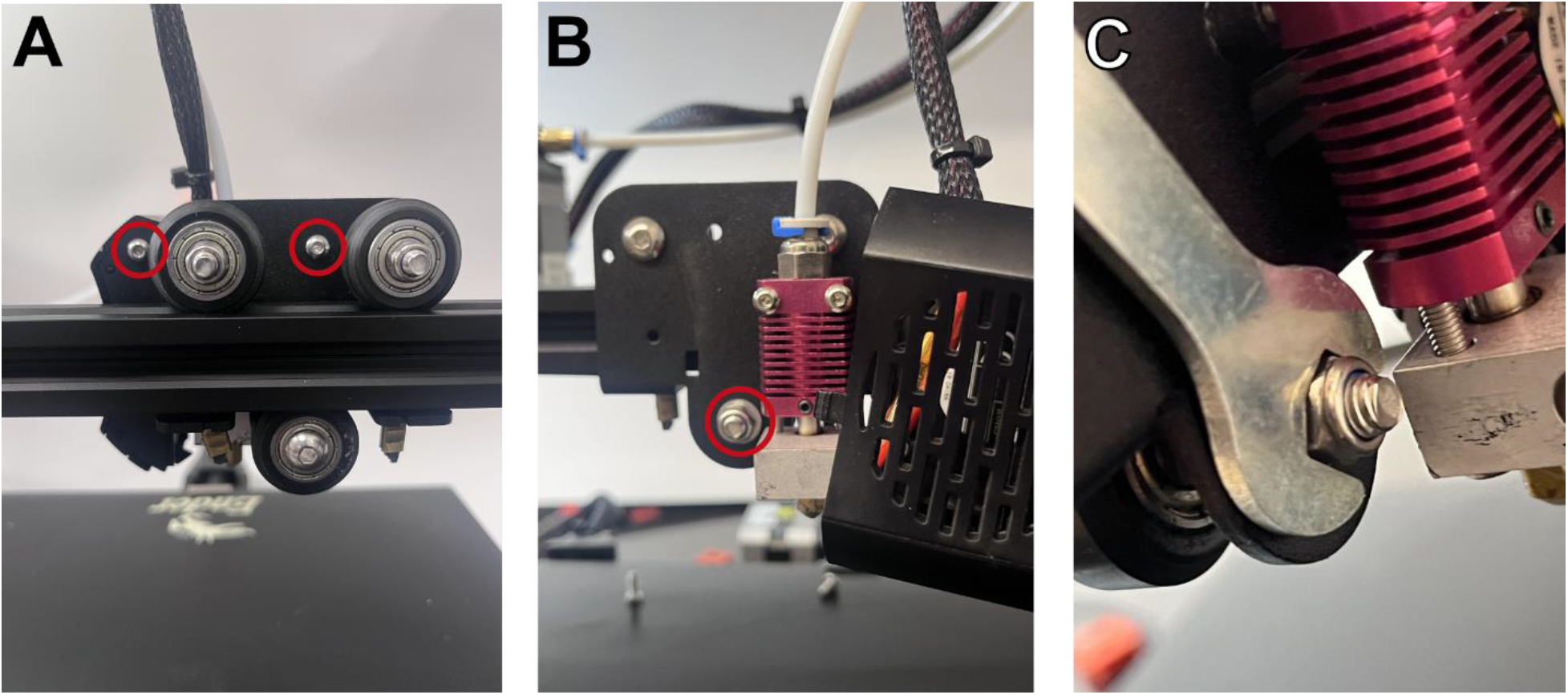
Removing the standard Ender extruder head. **(A)** The two screws that fasten the shroud to the extruder head are highlighted in red. **(B)** The locknut that must be loosened to loosen the bottom translating wheel of the carriage is highlighted in red. **(C)** Accessing this locknut with the included crescent wrench.

**Figure 4:**
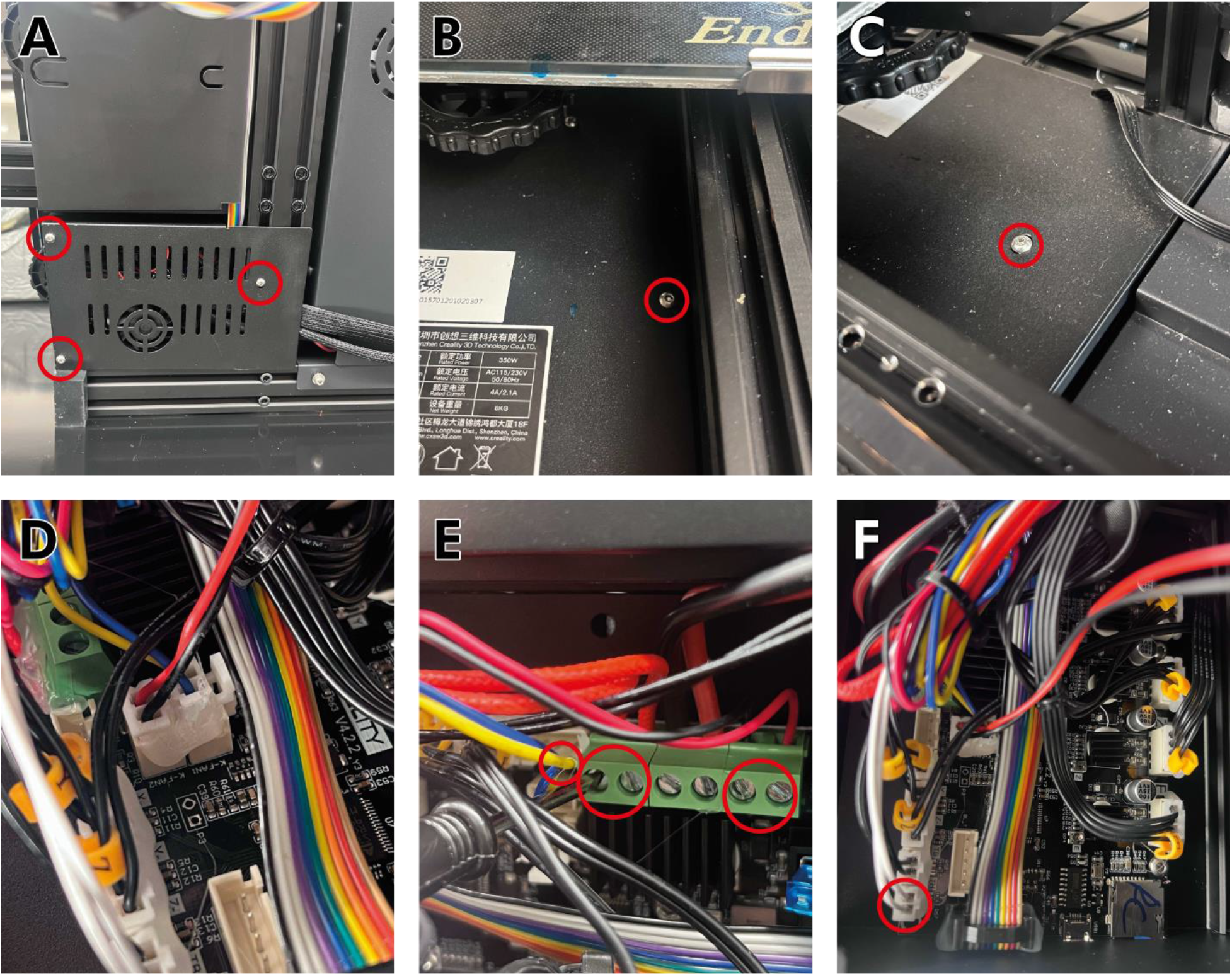
Removing the wires for the fans and hotend of the standard extruder. **(A)** Three screws at the bottom of the printer and two more screws underneath the build surface **(B,C)** must be removed to access the machine’s electrical wiring. **(D)** The cooling fan connector should be unplugged. **(E,F)** Wires attached to the extruder head (highlighted in red) may be pulled out or loosened with a flathead screwdriver.

### Constructing the Enderstruder

These steps assume you have already 3D-printed the five components in the Design Files. You can print these components on the Ender series printer you plan to modify. We can also print and ship these 3D-printed parts free of charge upon reasonable request.

1. Before beginning assembly, ensure you have gathered all the necessary 3D-printed and purchased components from the Bill of Materials. **Figure 5** shows all the materials necessary to build the Enderstruder.
2. Glue a hex nut into the slots on the top face of each of the 3D-printed herringbone gears. Make sure each hexagonal M3 nut is positioned with the corner facing downwards, as depicted in **Figure 6**.
3. Since the rolling carriage was removed from the v-slot extrusion in the previous set of instructions, we will need to install a new motion system. Here, we will replace the rolling carriage with a linear rail. To start, slide the MGN12H rail carriage onto the MGN12 linear rail. Then, place M3x6 screws on the 3^rd^, 7^th^, and 10^th^ holes, as depicted in **Figure 7A**. Slightly twist 3 T-Nuts into the bottom of the screws, and make sure the T-Nuts are aligned with the slot in the v-extrusion (**Figure 7B**). Place the linear rail on the Ender rail and tighten the screws with a hex key (**Figure 7C**). Ensure the rail is secure.
4. The 3D-printed carriage has a round and hexagonal hole. In the round holes, insert the ½" flanged bearing (it may need significant pressure to fit). Insert an M8 brass nut in the hexagonal hole. Align the carriage with the holes toward the bottom of the core, as shown in **Figure 8A**. Feed an M8x150 socket head screw into the bottom of the core and twist through the carriage until the screw pushes through the top of the core (**Figure 8B**). Make sure the carriage slit is facing toward the front of the mount and that it is suspended, as depicted in **Figure 8B**.
5. Next, the partially assembled core must be attached to the MGN12 carriage. Slide the mount until the four holes in the core shown in **Figure 9A** are aligned with the linear carriage. Fasten the core and carriage with four M3x16 screws (**Figure 9B**). WARNING: The M8x150 socket head screw may dislodge during this process.
6. Remove the stepper motor that is attached to the Bowden extruder, a common feature of Ender series printers. Position the motor beneath the large hole on the bottom of the core (**Figure 10A**). To avoid future wire entanglement, ensure that the pin connector faces the machine’s left. Fasten the core to the NEMA 17 stepper motor with four M4x16 screws until hand-tight (**Figure 10B**). WARNING: Over-tightening the screws can crack the 3D-printed part!
7. Place the M8 flanged bearings on the holes at the core (**Figure 11A**). Then, insert the smooth cylindrical rod through the diamond-shaped slot in the core and syringe carriage, as depicted in (**Figure 11B**). Apply downward force until the rod contacts the bottom of the diamond-shaped impression in the core.
8. Place the flexible shaft coupling on the motor’s drive shaft (**Figure 12A**). Do NOT tighten yet. Then, feed the partially threaded socket screw through the small herringbone gear and coupler, as depicted in **Figure 12B**. Tighten the included coupler set screws into the threaded rod and the drive shaft until the socket screw and drive shaft turn in unison (**Figure 12C, D**).
9. Slide the large herringbone gear on the threaded rod exposed at the top of the core (**Figure 13A**). Make sure that the herringbone gear teeth are correctly aligned. Fasten the gears to the threaded rods with M3x6 screws (**Figure 13B**). Tighten evenly, or the gear may sit at an angle. WARNING: Aligning the two gears may take some force. Be careful not to damage the teeth of the gear.
10. Place a loaded 10 mL BD syringe into the cavity on the core (**Figure 14A**). Align the syringe plunger with the carriage; this may require moving the carriage up and down by turning the large herringbone gear clockwise or counterclockwise until the top of the syringe is aligned with the carriage (**Figure 14B**). The Enderstruder is now complete! (**Figure 14C**).

**Figure 5:**
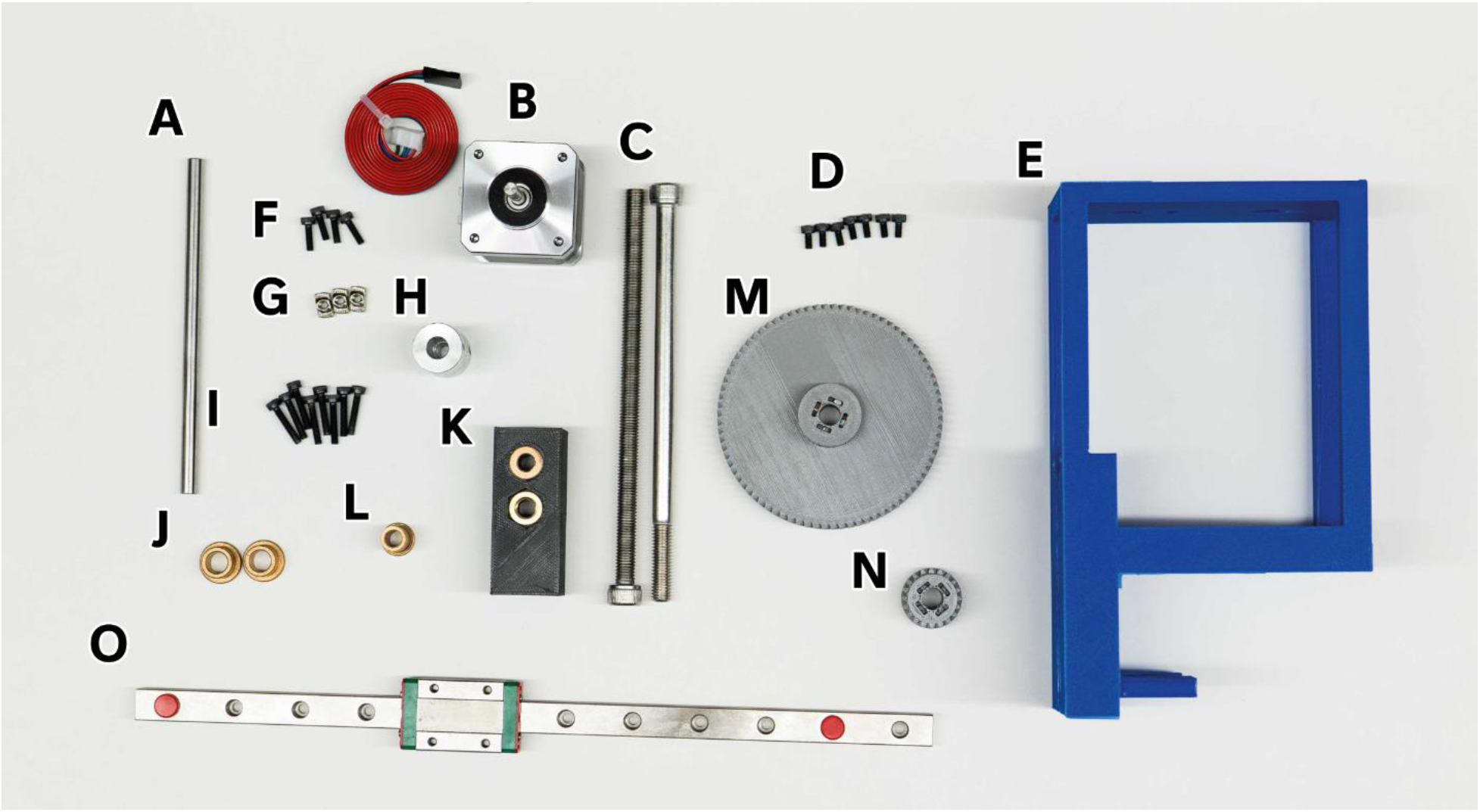
All 3D-printed and purchased components necessary for the assembly of the Enderstruder. **(A)** Linear shaft **(B)** NEMA 17 stepper motor **(C)**Threaded and partially threaded M8x150 socket screw **(D)** 8 M3x6 screws **(E)** 3D printed core **(F)** 4 M3X10 screws **(G)** T nuts **(H)** Coupler **(I)** 8 M3x16 screws **(J)** M8 Flanged bearings **(K)** Carriage with embedded coupling and nut **(L)** ½” Flanged bearing **(M)** Large herringbone gear **(N)** Small herringbone gear **(O)** Linear rail and carriage

**Figure 6:**
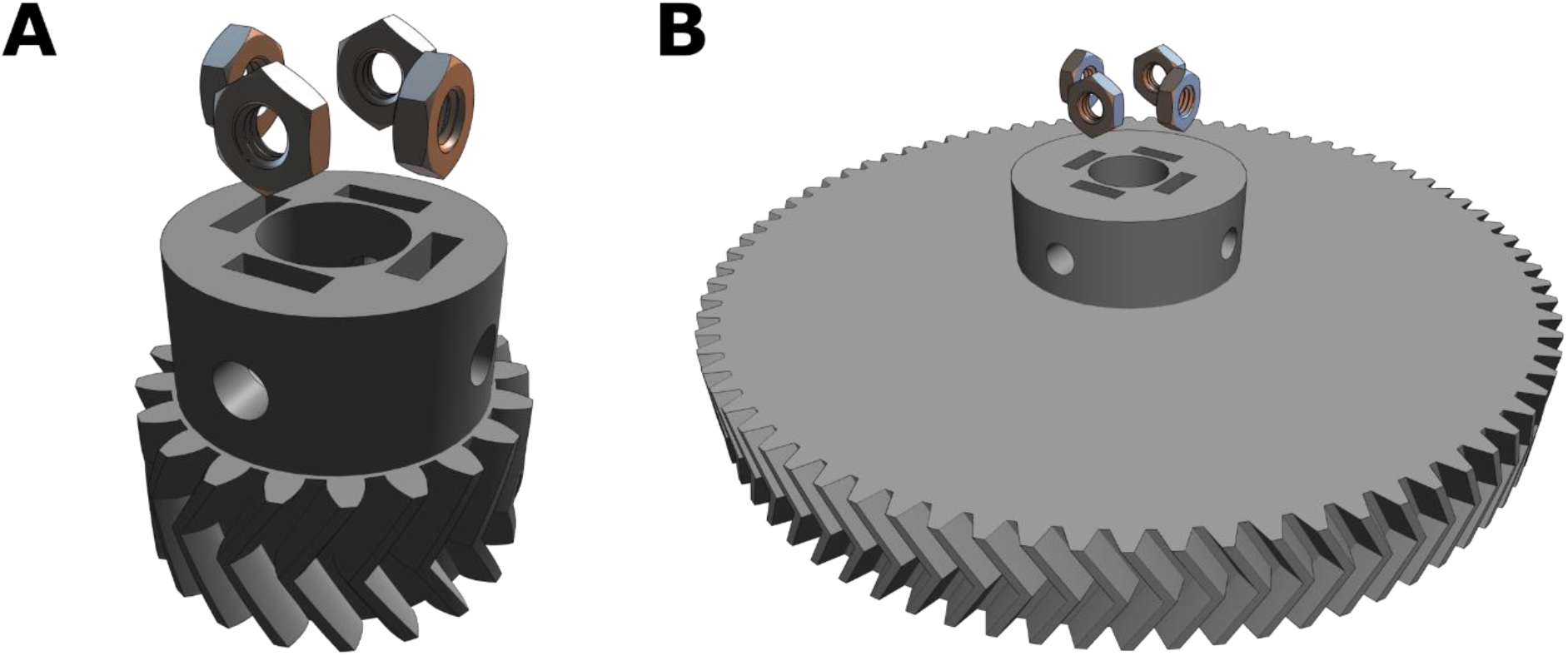
Gluing the hex nuts into the slots at the top of the two herringbone gears. Ensure that you complete this step first to ensure that the glue has time to bind and dry.

**Figure 7:**
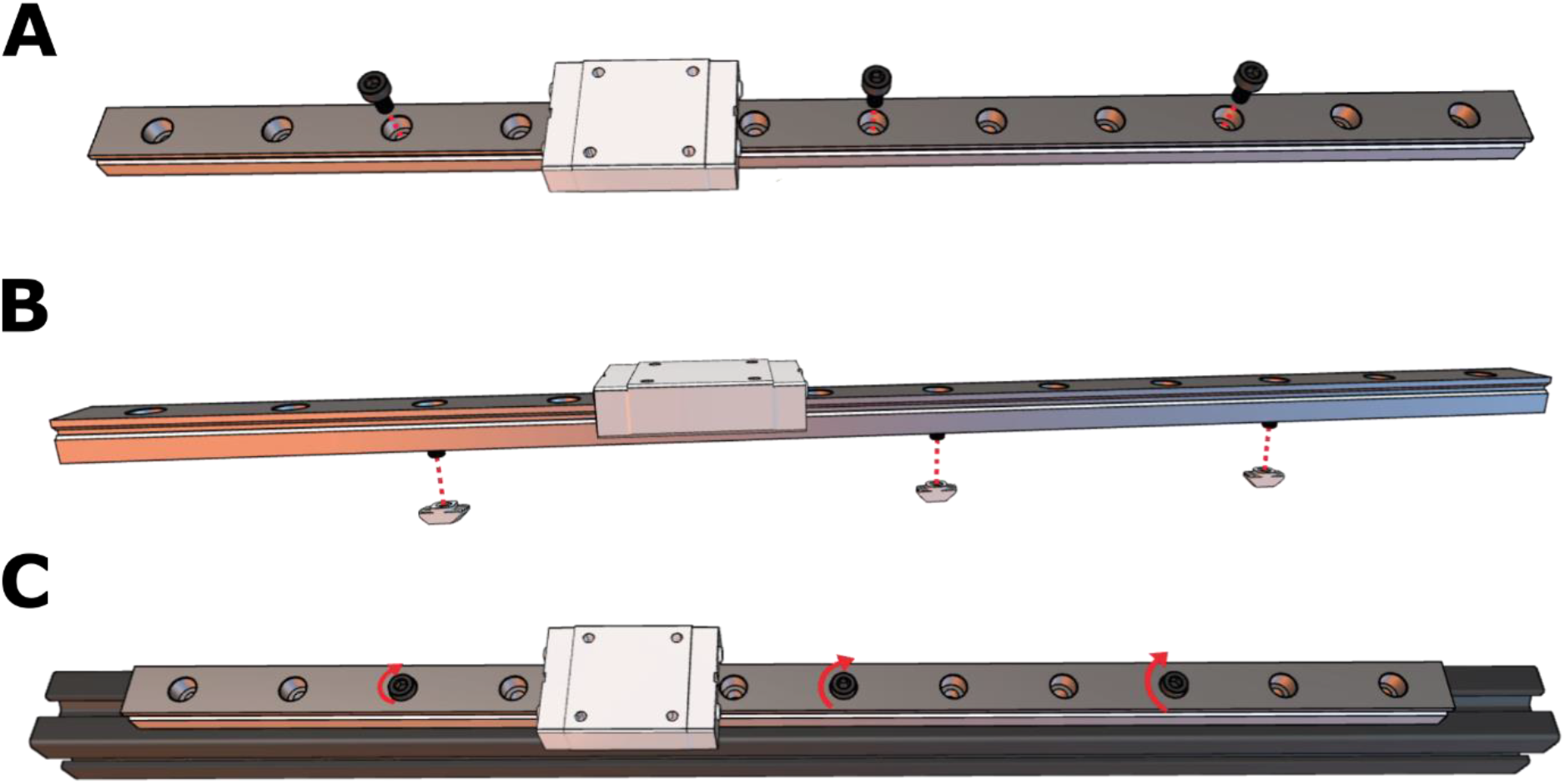
Replacing the rolling carriage with a linear rail. **(A)** Three M3x6 screws are evenly spaced along the rail. **(B)** T-nut are attached via a half-turn to the bottom of the screws and then **(C)** the rail is fastened to the extrusion gantry.

**Figure 8:**
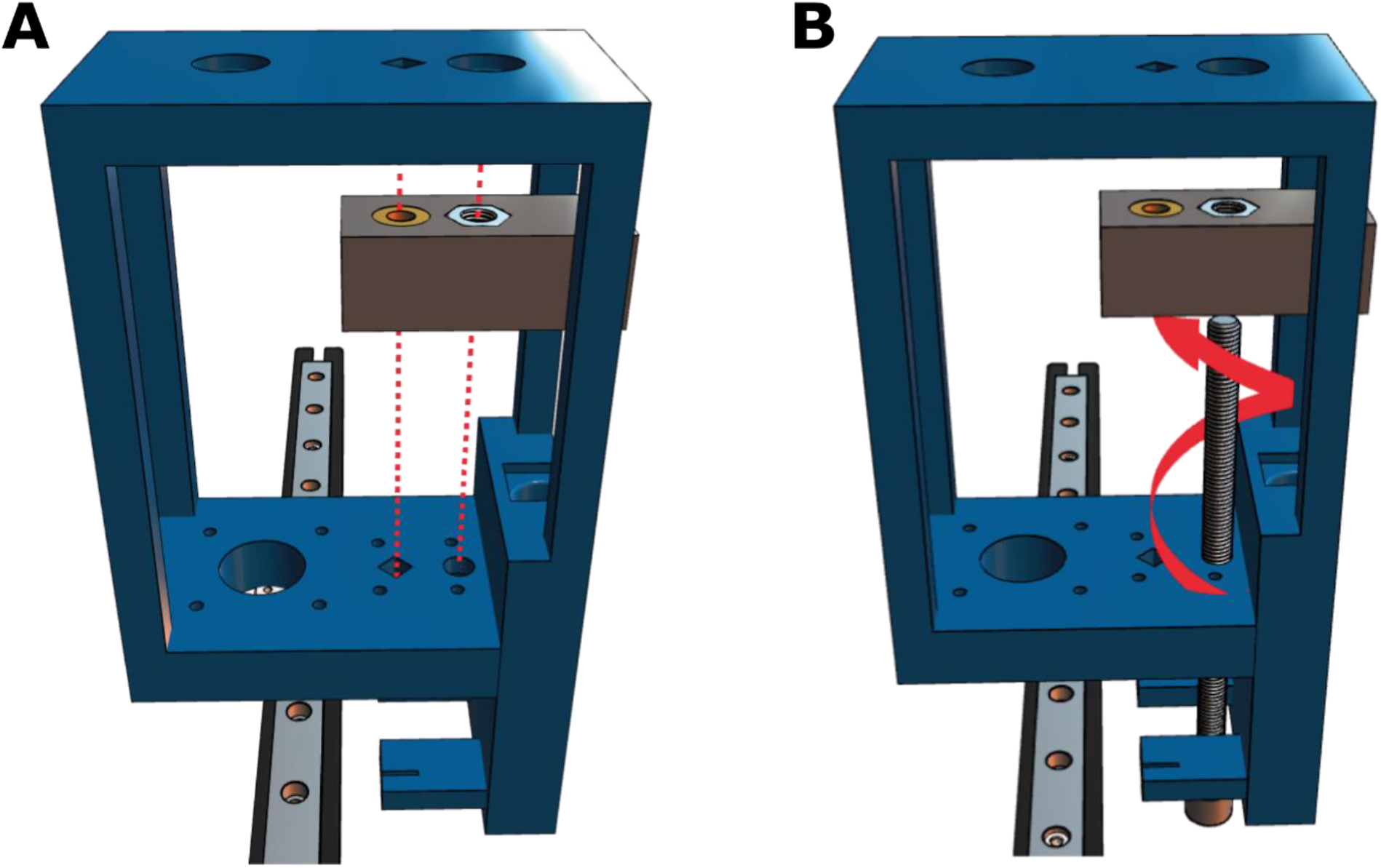
Threading the leadscrew through the carriage. **(A)** The carriage is positioned toward the front of the core and **(B)** a M8X150 socket screw is then threaded upwards and then fed through the top hole of the core.

**Figure 9:**
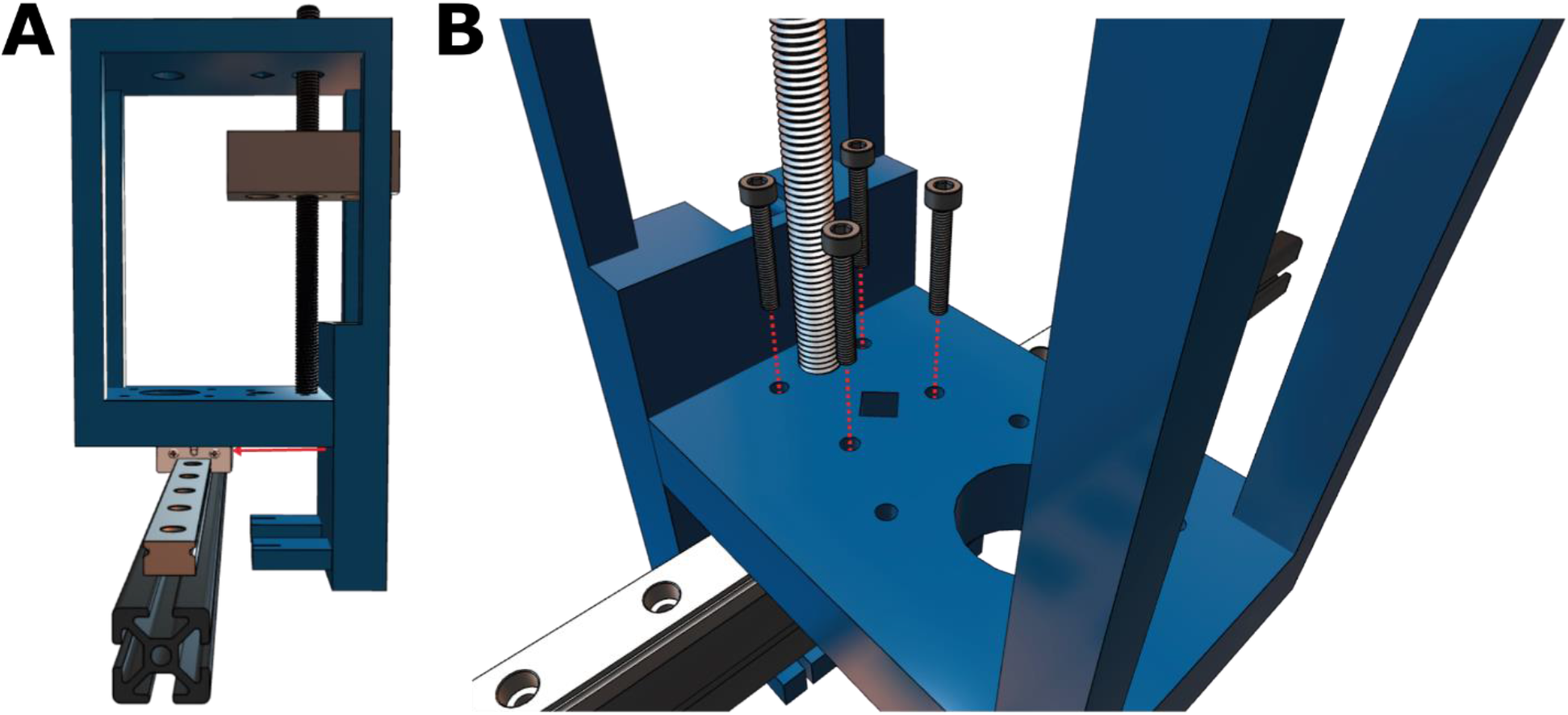
Attaching the core to the carriage on the linear rail. **(A)** The core slides over the linear rail and **(B)** is secured with four M3x16 screws.

**Figure 10:**
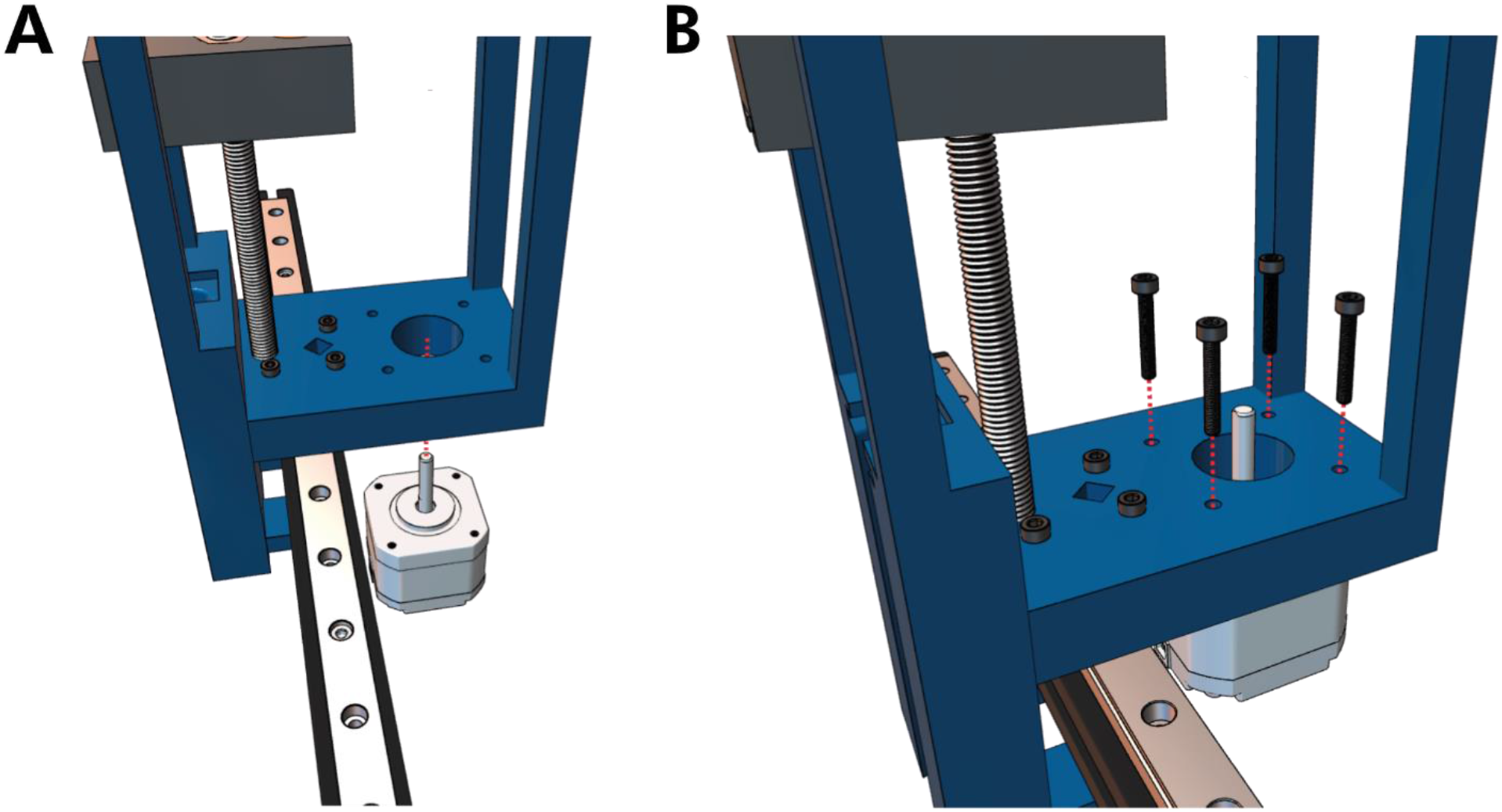
Attaching the NEMA 17 stepper motor to the Core. **(A)** The stepper motor is fed through the bottom of the core **(B)** and attached via four M4x16 screws.

**Figure 11:**
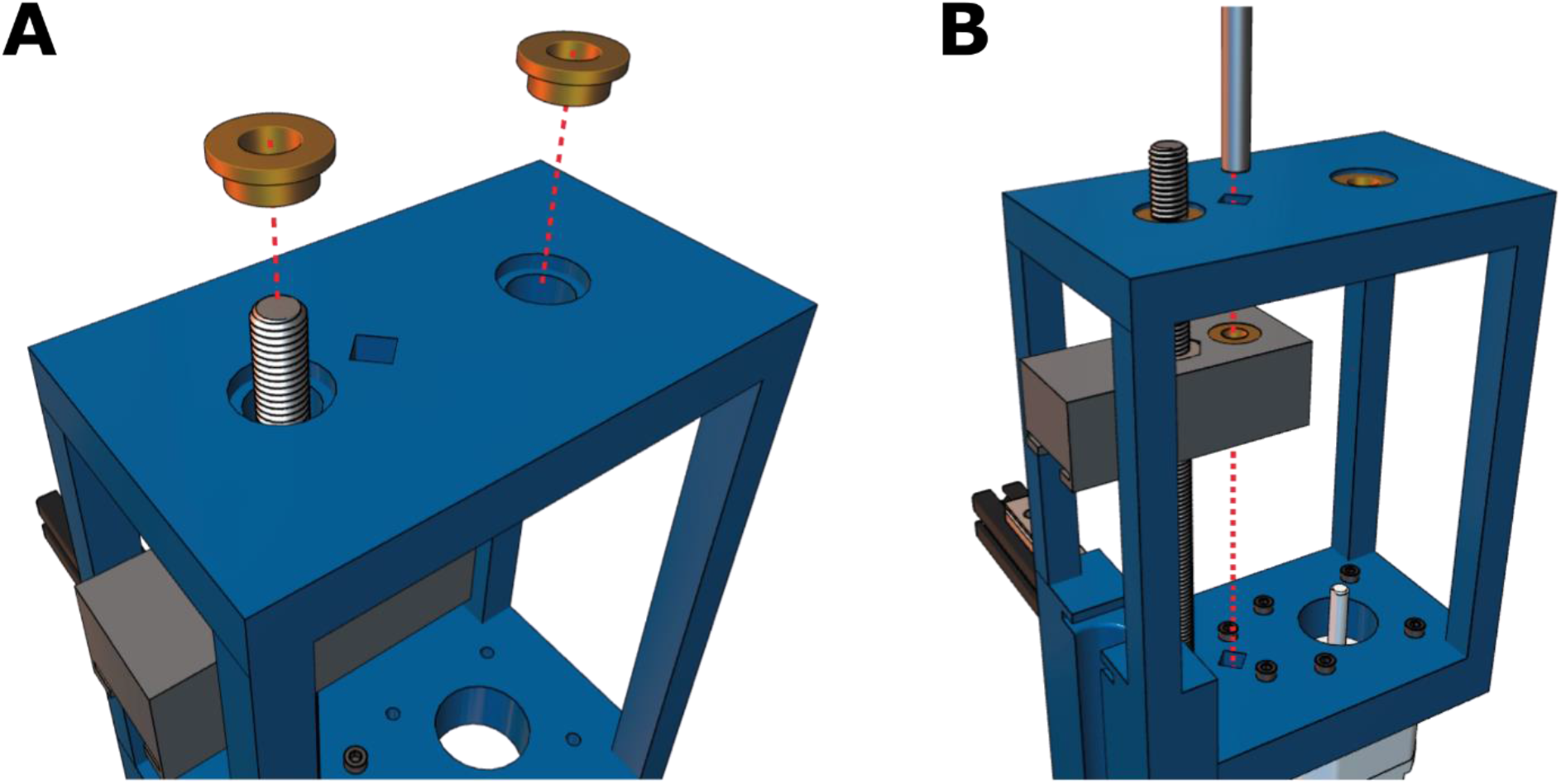
Adding the flanged bearings and cylindrical rod **(A)** Flanged bearings are inserted into the two large holes at the top of the core and **(B)** the cylindrical rod is fed through the diagonal hole of the top of the core and the rear hole in the carriage.

**Figure 12:**
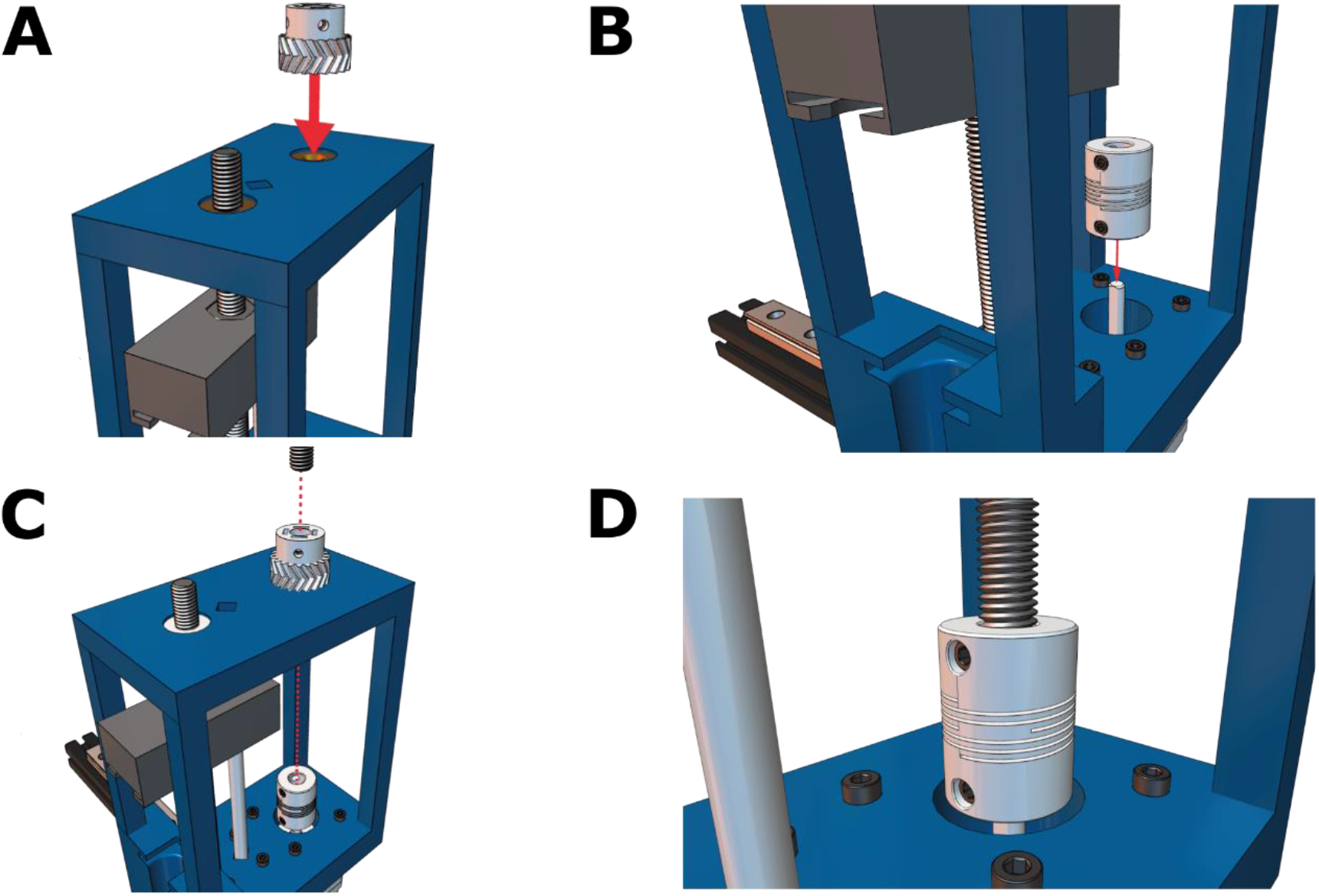
Coupling the leadscrew to the drive shaft of the stepper motor. **(A)** The small herringbone gear and the **(B)** flexible shaft coupling is positioned on the drive shaft of the stepper motor and then **(C)** connected via a partially threaded M8x150 socket headed screw. **(D)** The coupler set screws are tightened.

**Figure 13:**
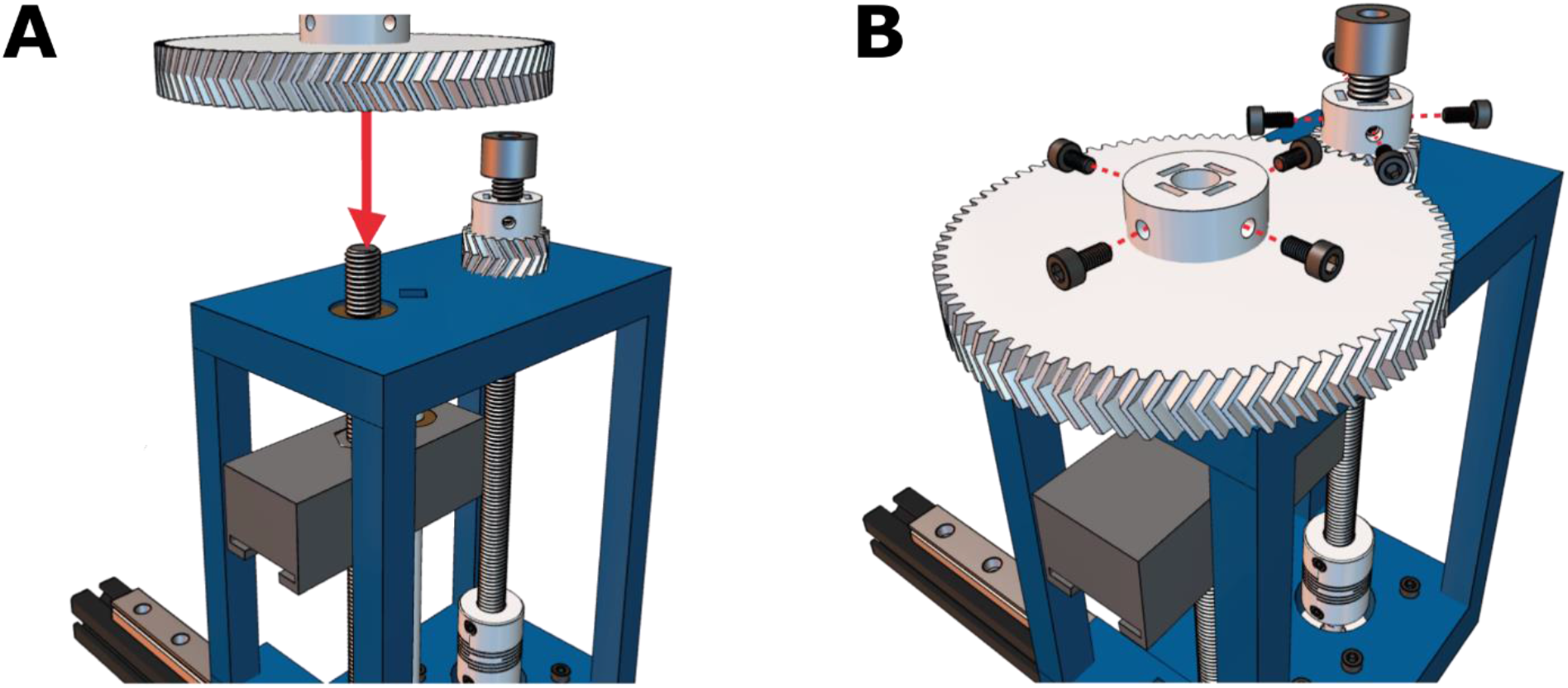
Adding the herringbone gears to the two socket head screws. **(A)** The larger gear is added second and **(B)** both gears are fastened through the hex nuts that were previously glued in Step 1.

**Figure 14:**
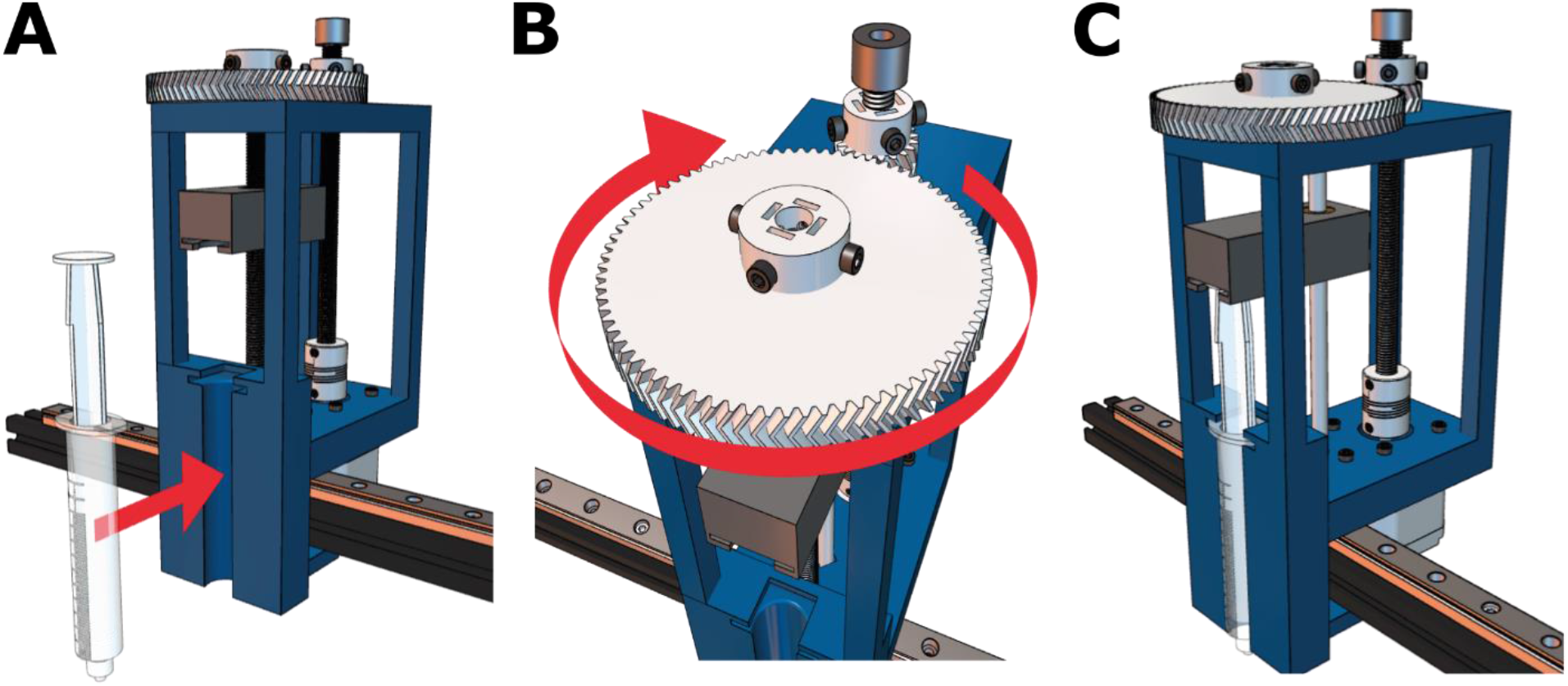
Loading and testing the Enderstruder. **(A)** The 10 mL BD syringe is loaded into the front slot, and **(B)** the large gear may need to be turned to align the carriage with the top of the syringe. **(C)** The final Enderstruder is shown mounted to the linear rail.

## 6. Operation instructions

Once the Enderstruder has been built, four additional steps must be performed before starting your first print. First, you will change the position of the x-axis limit switch. Second, the direction of the extrusion must be flipped; the default extrusion settings will result in retraction. Third, the E-steps will need to be calibrated. Fourth, UltiMaker Cura^TM^, our preferred open-source slicer, must be configured for a custom printer. At this point, the Enderstruder is ready for its first print with well-characterized material. We also outline how to create a printing profile for any biomaterial ink that is extrudable.

### Step 1: Changing the position of the x-axis limit switch

1. Since the Enderstruder has a significantly different form factor than the default extruder carriage on Ender series printers, it will collide with the z-axis metal bracket before contacting the x-axis limit switch mounted to the extruder motor assembly. (**Figure 15A**). Therefore, we printed an x-axis limit switch mounting block from Thingiverse (designed by user eschmidt12) [25]. This STL can be downloaded from Thingiverse for free (see reference for link).
2. First, remove the long screws from the plastic assembly that hold the extruder motor in place (**Figure 15B**). Then, use the two M3x6 screws that previously held the limit switch and fasten the switch to the 3D-printed piece (**Figure 15C**).
3. A M4x8 screw paired with a T-nut can then attach the assembly to the gantry. Ensure that the Enderstruder bumps the limit switch BEFORE contacting the metal bracket.

### Step 2A (Quick): Flip the polarity of the extruder motor plug

1. As in **Figure 4**, remove the screws from the bottom of the machine to access the mainboard.
2. You will see a group of 4 black wires with a yellow plastic tag with a black E printed on it and a black E printed on a white background (**Figure 16A**). To flip the plug, you must use the flush cutters to remove some plastic from the connection on the mainboard (**Figure 16B**).
3. Now, rotate the plug 180 degrees and replace the screws. The polarity of the motor should now be reversed.

### Step 2B (Preferred): Updating the firmware to reverse the polarity on the extruder

1. While Step 2A is straightforward from an implantation standpoint and is more accessible for inexperienced users, we prefer to edit the firmware to reverse the polarity of the motor. However, the latest firmware update will only work on the more recent 32-bit Ender series 3D printers. If your Ender has a mini-USB connection, it is an 8-bit machine, and only step 2A will work. If your Ender has a micro-USB connection, it is a 32-bit machine, and you can proceed below. The base firmware that the Enderstruder runs on is the Jyers fork of Marlin version 2.0.1. The Open Science Framework (OSF) repository linked in Specifications Table includes two firmware versions of the Jyers fork, one with the extruder motor reversed and the other with a standard extruder direction. Ensure that you have the correct firmware version uploaded. We have provided both if you would like to reverse the stepper motor in a future update. WARNING: Before proceeding, ensure the printer is switched off and the power is unplugged.
2. To do this, download the inverted firmware.bin file found on the OSF depository and upload this .bin file to an empty Secure Digital (SD) card. Please note that this card must have at least 8 GB of free space, should be in FAT32 format, and must have an allocation size of 4096.
3. Rename the file to Firmware001.bin. WARNING: If you perform future updates, rename the file on every firmware update attempt.
4. Insert the SD card containing the file into the printer’s SD card slot.
5. Turn the printer back on and wait for the printer screen to turn on and load fully. Once the screen has loaded, turn off the printer and remove the SD card. Remove the firmware from the card’s memory if you want to use this SD card for future G-code uploads. The polarity on the stepper motor should now be reversed.
6. Next, you will need to upload the new touchscreen files accompanying the Jyers fork. These files can also be found on the OSF depository. Ensure that the printer is turned off and disconnected from power, and remove the touch screen from the printer.
7. We will need to use a microSD card to update the touch screen. Remove the four screws from the back of the touchscreen display to reveal the PCB (**Figure 17A**).
8. Take an empty microSD card and format it in FAT32 at a 4096 allocation size. Move the firmware files on the OSF repository onto the empty SD card and then safely eject it.
9. Place the SD card into the SD card slot in the back of the touchscreen, as shown in **Figure 17B**. Connect the touchscreen back into the printer using the rainbow connector and turn on the printer. The touchscreen should flash to a blue screen with orange text (**Figure 17C**).
10. Once the screen says update finished or the screen flashes to reading "Creality," turn off the printer and remove the SD card from the touchscreen. The touchscreen update is now complete.

### Step 3: Calibrating the E-steps of the Enderstruder

The number of discrete angular displacements (extruder steps, or E-steps) a stepper motor uses to extrude 1 mm of a plastic filament is a critical setting for FFF machines. We used the original NEMA 17 stepper motor included with the Ender series model; therefore, our motor has 200 steps per revolution (1.8 degrees per step).

**Figure 15:**
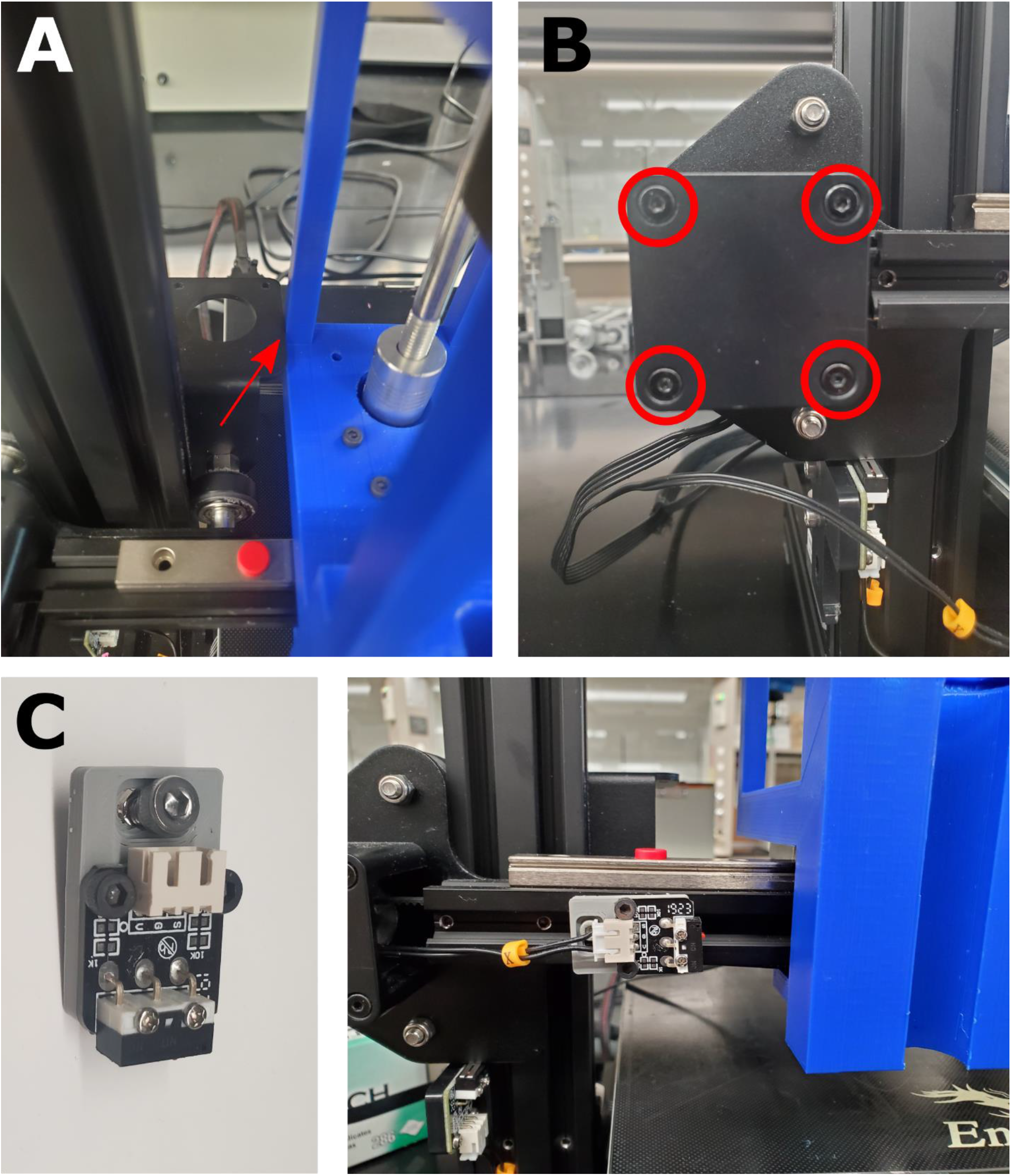
Moving the x-axis limit switch avoid a collision. **(A)** If the x-axis limit switch is not moved, the collision marked with the red arrow could occur. **(B)** Remove the long screws to access the limit switch and unscrew it from the plastic base. **(C)** Use these screws to fasten the limit switch to the 3D-printed piece and gantry (right) so that the Enderstruder encounters the limit switch before damage occurs when homing the syringe extruder.

**Figure 16:**
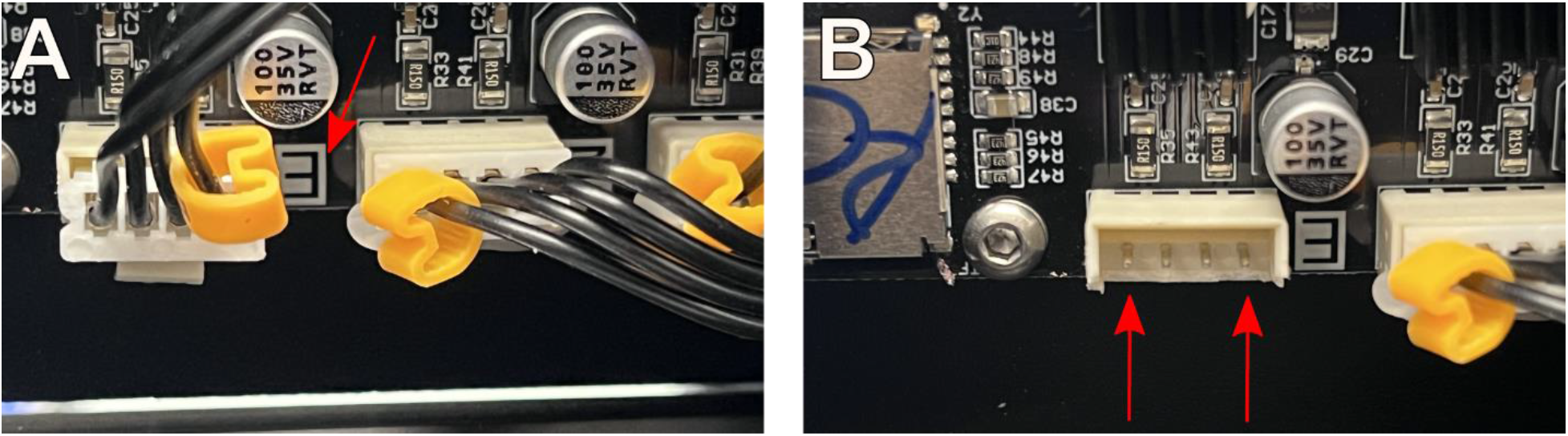
Flipping the polarity of the extruder motor. **(A)** Remove the screws on the bottom of the machine to access the mainboard. Identify the plug that belongs to the extruder motor. **(B)** Clip with flush cutters as necessary to allow the plug to be flipped.

**Figure 17:**
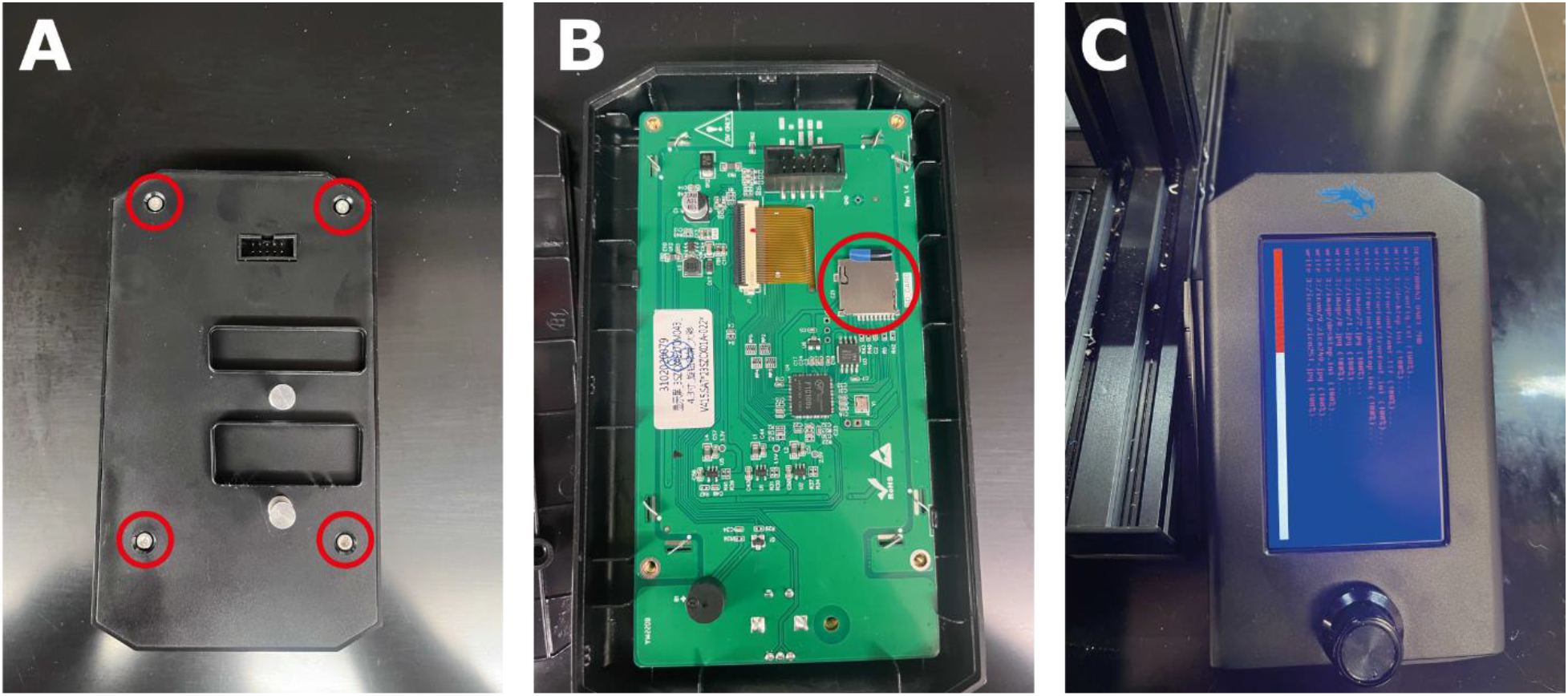
Flashing new firmware to the Ender machine. **(A)** The SD card slot **(B)** is accessed by removing the screws circled in red. **(C)** The loading screen that should appear during this process.

While FFF 3D printers may require an average of 100-200 E-steps to print PLA or similar thermoplastic materials, bioprinters may require thousands of E-steps to achieve similar extrusion rates. This difference is due to the difference in extrusion mechanisms. The E-steps for an Ender series thermoplastic 3D printer equipped with a Bowden extruder and 10.9 extruder gear can be calculated as follows:

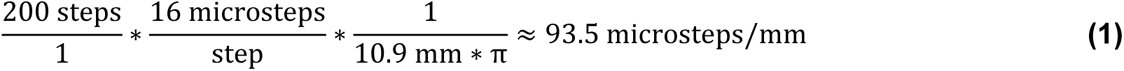

The user may notice that their stock Ender machine often comes pre-loaded with an E-step number in this range. In contrast, most syringe extruders drive a threaded nut along a leadscrew to depress the syringe’s plunger. Therefore, the number of revolutions of the stepper motor must be matched to a linear vertical displacement. These calculations often result in considerably larger E-step values. For example, open-source designs by Dávila *et al.* [17] and Bessler *et al.* [13] require 1600 E-steps/mm and 2560 E-steps/mm, respectively, with 200 steps per revolution (1.8 degrees) stepper motors. Like Bessler *et al.*, we use a M8 screw with a pitch of 1.25 mm; however, we also added a gearing ratio of 4:1 to increase the torque generated by the stepper motor. Therefore, our theoretical E-steps can be calculated as follows:

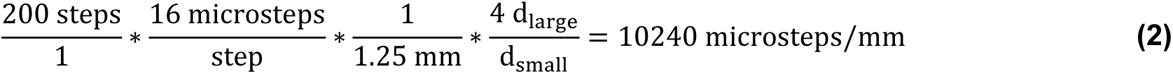

Bioprinted scaffolds are typically much smaller than models printed on FFF printers; therefore, over-extrusion and under-extrusion caused by improper E-step calibration will be much more noticeable. Like thermoplastic printers, syringe extruders often exhibit E-step values different from theoretically calculated ones. Below, we outline how to conduct an experimental E-step calibration that considers slippage, back- pressure, and other non-calculated factors.

1. Download the Pronterface program from the website https://www.pronterface.com/. Connect your Ender series 3D printer with a mounted Enderstruder to Pronterface by connecting it to your computer using a micro-USB cable.
2. Load an empty 10 mL BD syringe with roughly 5 mL of water while minimizing the air bubbles. Slot the loaded syringe into the Enderstruder. You may need to manually move the gears to get the carriage piece in the proper position. Find an empty container (e.g., a small glass beaker) and tare it on an analytical balance.
3. To enable cold extrusion, enter the following G-code into the terminal: M302 S0. Set the extrusion distance to 3 mm and the extrusion speed to 1 mm/min.
4. Next, enter M503 into the terminal. This command will display a list of all the current settings stored on the printer’s mainboard. Look for the line of code that contains "steps per unit." The code should look like M92 X80.00 Y80.00 Z422.61 E**93** (93, the stock number of E-steps).
5. To begin, set E to the theoretical value: 10240. This E-steps value will change if you modify the leadscrew or gearing ratio.
6. Raise the Enderstruder and place your empty container directly under the syringe. Then click the "Extrude" button, and the Enderstuder will move the syringe plunger to what the machine thinks is 3 mm. Remove any residual drops of water after the plunger has completed its movement with a Kimwipe or similar absorbent material.
7. Place the empty container back on the scale and record the increase in mass from the extruded water. We can calculate the new E-steps value from equations (3) and (4):

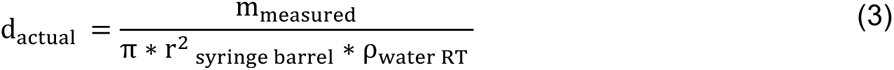

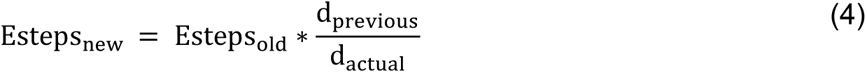

d_actual_ is the distance traversed by the plunger in the z direction in millimeters; m_measured_ is the mass of the water extruded, r_syringe barrel_ is the radius of the syringe barrel (7.25 mm for a 10 mL BD syringe), ρ_water RT_ is the density of water at 23° C (we used 997.5 kg/m^3^), d_previous_ in this example is 3 mm, Esteps_old_ in this example is 10240, and Esteps_new_ is the calibrated E-step value.

8. In our experience, obtaining an Esteps_new value that moves the plunger to the expected distance may take three to five consecutive trials. We have uploaded sample data to the OSF repository as an Excel table for reference.

### Step 4: Installation and configuration of Ultimaker Cura^TM^

We prefer to use Ultimaker Cura^TM^, an open-source slicing software. This section details how to download and optimize the software for the Enderstruder.

The most up-to-date version of Cura^TM^ (must be v5 or later) can be downloaded using the following link: https://ultimaker.com/software/ultimaker-cura. Follow the installation wizard to set up Cura^TM^ on your computer, and then follow the steps below to set up the printer profile for the Enderstruder.

1. Once prompted with the "Add a Printer" dialogue box, select "Add a non-networked printer." Scroll down the drop-down menu and select "Custom è Custom FFF." Rename the printer "Enderstruder" and then click "Add the printer."
2. Change the printer settings to match the highlighted areas under the **Printer** (**Figure 18A)** and **Extruder 1** (**Figure 18B**) tabs. Ensure that your nozzle diameter matches that of the needle that is being used. For example, a 27-gauge needle has a diameter of 0.21 mm. Other needle gauges can be easily found via any search engine.
3. Paste the following G-code into the Start Gcode box. Please note that anything following a semicolon is a comment not read by the compiler. **M302 S0**; Enable cold extrusion **G92 X0 Y0 Z0.00 E0.00**; Tell the printer that the nozzle is homed and at a zero Z height to enable printing to begin at the manually-selected point. The X and Y values used correspond to one-half the length of the printer platform’s X and Y build dimensions. **T0**; Set the nozzle temperature to zero
4. Paste the following G-code into the End Gcode box: **M104 S0**; Turn off temperature **M106 S0**; Turn off the fan **G92 E0**; Set extruder value back to 0 **G92 Z0;** **G1 X20 Y20 Z20**; **M84**; Disable motor
5. Download the BioprinterMaterial.xml.fdm material file from the OSF repository. In Cura^TM^, select "Configure settings" at the top left-hand corner under the Settings tab. Once the Preferences box appears in the list to the left of the box, select "Materials." Then, select "Import" and choose the bioprinting material profile you downloaded, named BioprinterMaterial.
6. We have created default printing profiles for five materials paired with the needle with the smallest diameter that the Enderstruder can reliably extrude at ambient temperature. Select "Profiles" instead of Materials and import the desired Profile into Cura^TM^.

**Figure 18:**
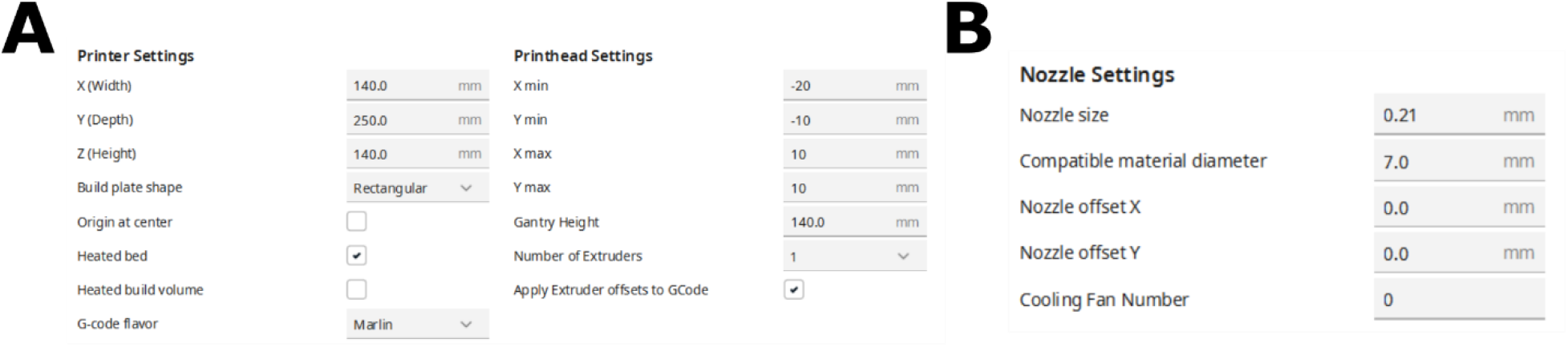
The settings for the **(A)** Printer and the **(B)** Extruder. They should appear as separate tabs in the same popup window.

### Step 5: Completing a print with a well-characterized material

The Enderstruder is now ready for printing! We typically print via a USB connection; most prints take less than an hour to complete.

1. Open Cura^TM^, and in the top left corner, go to the *Help* tab. Select *Show Configuration folder → Definitions Changes.* Look for a file named *Enderstruder_settings.inst.cfg.* If you are using a Windows machine, we suggest using Notepad to open and edit this file; on MacOS, we suggest using TextEdit.
2. There will be a line in this file that will read *machine_heated_bed = False* (**Figure 19A**). After this line of code, paste *machine_nozzle_temp_enabled = False.* Save this file, exit, and re-start Cura.

a. Note: If *machine_heated_bed = False* does not exist inside this file, insert it above *machine_start_gcode = …*
3. Upon re-starting, confirm that you have selected the correct print and material profiles. Click on the drop-down menu shown in **Figure 19B** to open the Print Settings. Select **Show Custom Settings.** If ever prompted to keep or discard, always select discard unless you want to change the printing profile permanently.
4. Load in the model that you would like to print using the folder icon in the top left corner. The model will need to be in stereolithography (STL) format. We suggest starting with a simple 3D geometry like a cube or cylinder. These can be imported via the "Part for Calibration" extension in Cura. Ensure the model is oriented in the direction you would like.
5. Drag the model to the bottom left-hand corner of the build plate, i.e., the origin, in Cura and press the Slice button. Click the preview button on the top and confirm that there are no errors (if the model does not appear as it should) in the sliced model.
6. We typically print on two surfaces: a spray-painted microscope slide or a trimmed weigh boat. We use these surfaces because they provide a waterproof barrier with a flat white finish that minimizes reflections when imaged on a stereomicroscope. We also print in suspension baths, including Freeform Reversible Embedding of Suspended Hydrogels (FRESHv2.0), pioneered by *Lee et al.* [26]. Before printing, it is essential to ensure the needle is at the correct height, as the start G-code will not change the z height. For a print in the air, we recommend placing the needle *just* above the printing surface (**Figure 20A, B**). In a suspension bath, the needle should be placed toward the front left of the container and toward the bottom (the z-distance is less critical) (**Figure 20C**).

a. Slightly lower the needle until it touches the bed. Once it touches slightly, move the bed back or forward toward your desired print area; the needle should deflect slightly during this bed movement. When this happens, twist the shaft coupler until the needle is again perfectly vertical.
7. Turning the large herringbone gear may be necessary to extrude some material and initiate flow. You can begin the print by clicking *Print via USB* in the bottom right of the screen.
8. When the print has been completed, the stepper motors should be disabled (M84), and the bed can be pulled toward the user. An error may occur, and the stepper motors will not be disabled. To do this manually, select Prepare → Disable Stepper on the Ender 3 V2 LCD screen.
9. It may be necessary to stop a failed print in a suspension bath. In Cura, press the "Pause" button and wait for the print to pause. Once paused, raise the z-axis by pressing the up arrow above the Z button until the needle clears whatever container is used to hold the FRESH.

**Figure 19:**
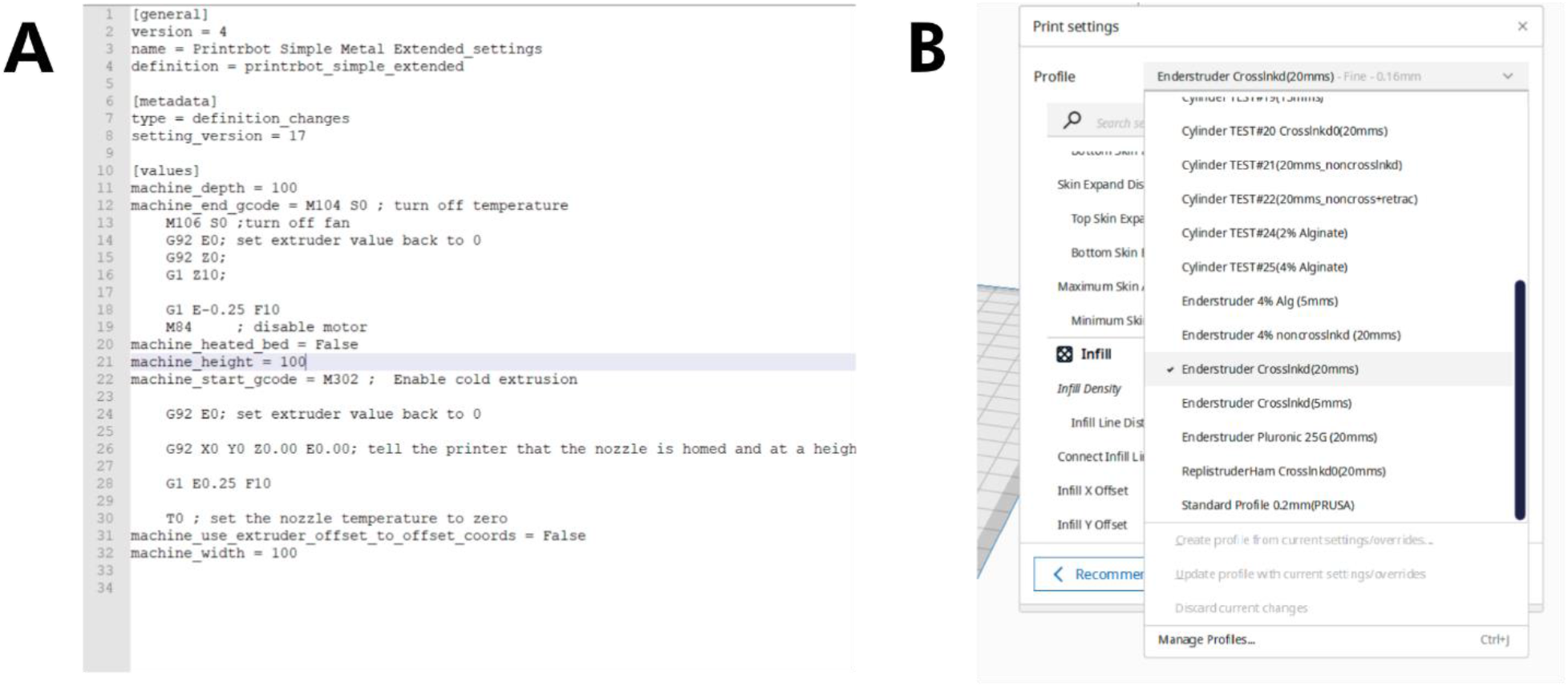
Changing printer settings. **(A)** Ensure that cold extrusion is enabled and **(B)** that you have the correct print profile selected.

**Figure 20:**
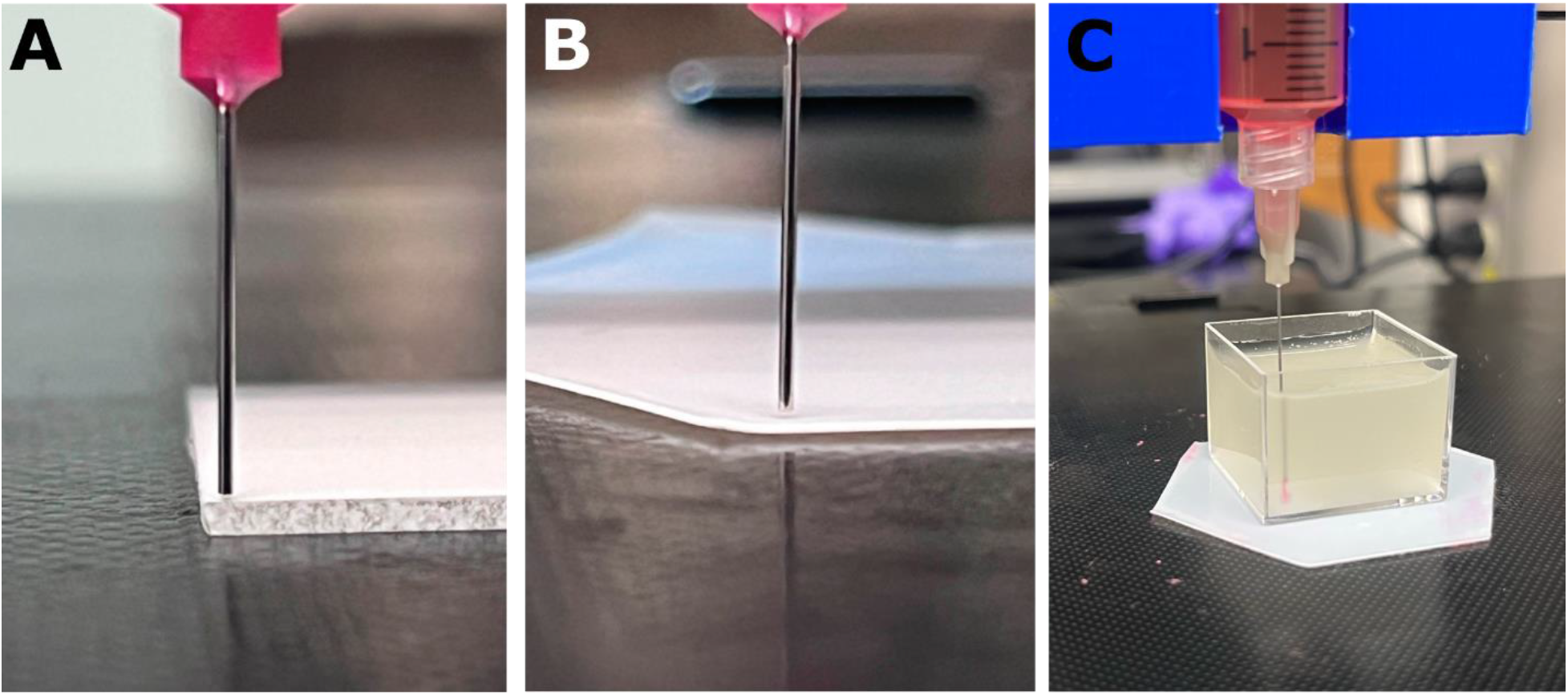
Correct nozzle height for **(A)** a glass slide, **(B)** a trimmed weigh boat, and **(C)** FRESH suspension bath.

### Step 5: Calibrating new print profiles for novel or poorly characterized materials

When making a new print profile, it is especially critical to tune the settings for the walls, the bottom/top layer, and the infill. The primary setting that will be used to adjust these features is material flow, which controls how much material is extruded out of the syringe. Below, we detail how to tune your printing profile by iteratively printing a simple calibration cube.

1. Go to the extension tab in Cura, choose "Part for Calibration," then "Add a cube." Scale down the cube to 10 mm and place it close to the front left corner of the bed. Ensure that "skirt distance" is set to 3 mm in the printer profile setting.
2. Set the wall count to 0, infill to 0, and bottom/top layer to 0. Start with the flow rate set to 100% and print.

a. Pause and then abort prints to save material once you have determined the quality of the print. To do this, click the "End Print" button while the Ender prints.
3. Repeat Step 2, but gradually decrease the flow rate in increments of 10% until the material does not extrude during printing. Once this is done, continue conducting print tests but increase in increments of 5% until the material starts to flow consistently out of the needle.
4. Once your flow setting is set to your desired value, repeat Steps 2 and 3 while varying the following settings: Wall Flow, Outer Wall Flow, Inner Wall Flow, Top/Bottom Flow, Infill Flow, Skirt Flow, Initial Layer Flow, and Infill Flow.

a. Change your line counts. For example, to test the impact of flow rate on the integrity of the bottom/top layer, set the wall line count to 0, infill to 0, and bottom layer to 1, thereby isolating that setting.

i. **<u>Walls</u>**: The cube’s walls, or sides, should be printed with no holes or areas of under/over extrusion. The flow setting that affects the walls is wall flow. This setting is changed across multiple test prints until the walls are correctly extruded with no holes or under/over extrusions (**Figure 21A**).
ii. **<u>Bottom/Top Layers</u>**: An ideal bottom and top layer will not have holes and be thin enough that it does not interfere with future movements of the needle (**Figure 21B**). The principal setting that determines the thickness of these layers is called bottom/top layer thickness and should be set at the inner diameter of the needle. The flow setting adjusted to calibrate the bottom and top layers was bottom layer flow and top layer flow, which refers to the flow setting that only affects the bottom layer and top layer of a print.
iii. **<u>Infill</u>**: The infill of a 3D print lends structural rigidity to the walls while conserving material (**Figure 21C**). The flow setting that is adjusted is the infill flow. The infill flow is adjusted until the infill is connected; in contrast to the walls and bottom/top layers, some over-extrusion should not affect the print quality.
5. Once all flow rates have been calibrated, increasing the print speed to 15-20 mm/s is usually desirable to reduce total printing time. You should increase the print speed by 5 mm/s increments from a base speed of 5 mm/s. Typically, this will result in a steady increase in the flow rates, i.e., adjusting the settings found in step 4.

**Figure 21:**
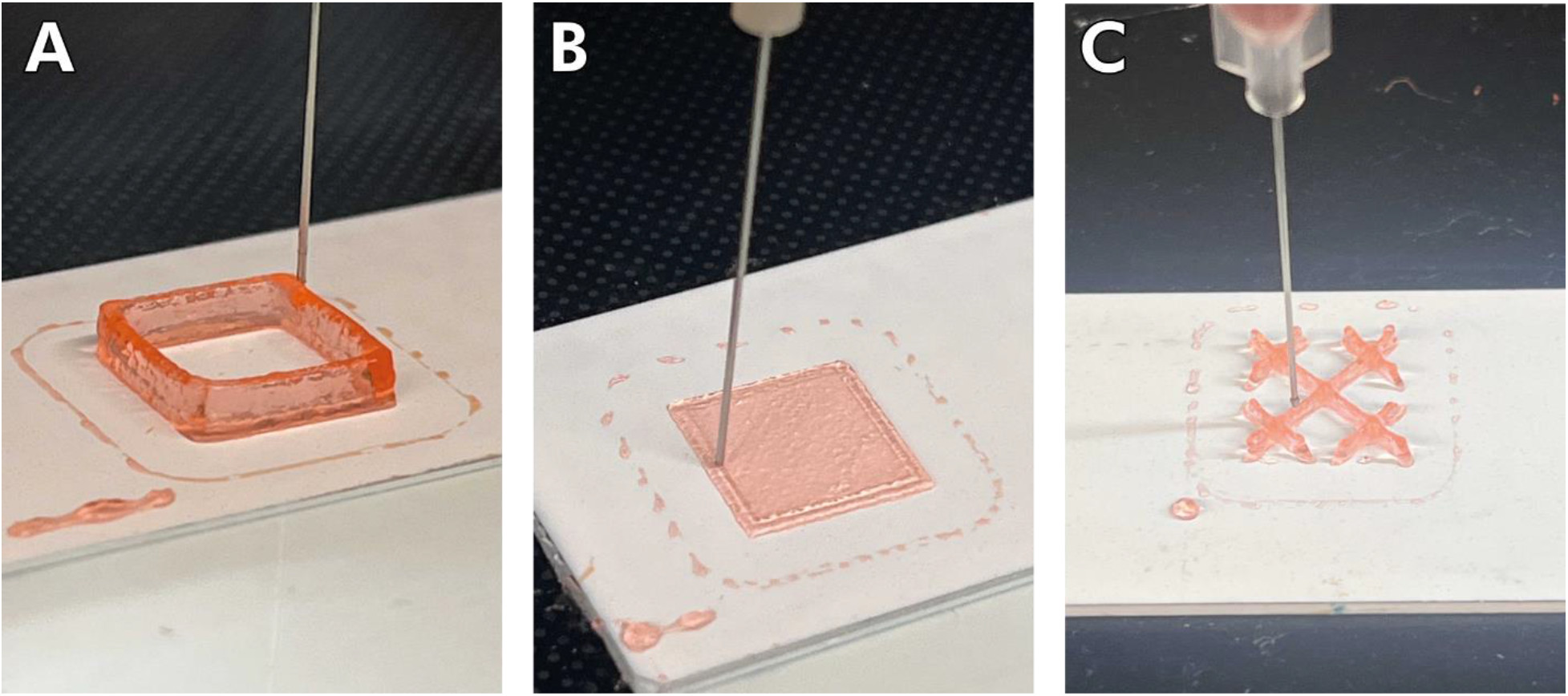
Sample geometries of 3.5% w/v CaCl_2_ cross-linked alginate printed during flow rate calibration. **(A)** Walls should be continuous and have relatively sharp corners, **(B)** layers should also be continuous and not extend beyond the square border and **(C)** grid infill should appear as shown above.

## 7. Validation and characterization

We performed standard evaluations of known biomaterial inks to complete an in-depth validation and characterization of the Enderstruder. We specifically tested five materials that we identified as some of the most used in extrusion bioprinting applications: 4% w/v alginate, 3.5% w/v CaCl2 cross-linked alginate, 10% w/v gelatin methacryloyl (GelMA), 40% w/v Pluronic F-127, and stock NIVEA Creme. Alginate and GelMA are biomacromolecules that can be ionically cross-linked and photo-cross-linked, respectively. Pluronic F-127 is a poloxamer that solidifies upon warming, and NIVEA Crème is a Bingham plastic (i.e., self-supporting fluid under a static load) that consists primarily of water, mineral oil, petrolatum, glycerin, and wax. We completed three tests for each material: filament uniformity, filament fusion, and printability. These printability tests and their origins have been covered in detail by Fu *et al.* and Schwab *et al.* [27,28]. Briefly, filament uniformity attempts to predict the final fidelity of the printed construct by comparing the width of extruded lines of filament; a high degree of uniformity is usually predictive of high fidelity. The filament fusion test attempts to estimate the minimum resolution that two adjacent filament strands can be placed without surface tension merging the two strands into a single entity [29]. This test is done by extruding a "zig-zag" pattern with decreasing distance between each set of strands [30]. We estimated printability using the scoring system first promulgated by Ouyang *et al.* [31]; briefly, we printed a 10 mm x 10 mm 3-layer woodpile lattice with 16 total pores and calculated the printability score (P) as a function of the circularity (C). Circularity is a parameter defined in the "Analyze Particles" plugin in ImageJ, where *A* represents the shape’s area, and *L* represents the shape’s circumference.

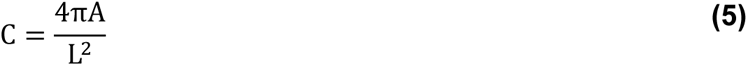

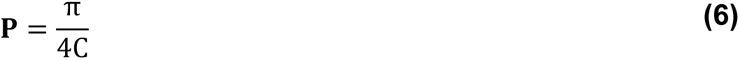

For all three tests, we imaged the resulting patterns on a binocular stereo microscope and then used a short custom ImageJ script that automated the process and minimized user bias. These ImageJ script is available on the OSF repository. We concluded this testing phase by printing challenging 3D geometries used to test traditional FFF printers’ capabilities.

Our findings showcased a high degree of filament uniformity for all tested biomaterial inks, with variations in line width within a tolerable range of 110 microns (**Figure 22**). Similarly, only GelMA and fluid alginate experienced a loss of resolution due to adjacent surface tension (**Figure 23**). Fluid alginate was too fluid to stabilize visible pores; however, cross-linked alginate, Pluronic F-127, GelMA, and Nivea Crème attained average printability scores of 1.30, 1.17, 1.33, and 1.05, respectively (**Figure 24**). Additionally, the Enderstruder could print complex designs with extended printing times and the need for retraction, exemplified by the successful printing of a twisted cube [32] and the ubiquitous "3D Benchy" calibration model (**Figure 25**). To further enhance the Enderstruder’s capabilities, we aim to implement temperature control, incorporate an onboard cross-linking mechanism, and devise a more compact form factor. Moreover, a detailed COMSOL analysis could observe and mitigate shear stress on printed materials, thereby enhancing cell viability. By presenting the Enderstruder and its iterative development process, this study will contribute to the growing repository of open-source bioprinting solutions, fostering greater accessibility and affordability for researchers in tissue engineering and other related disciplines.

**Figure 22:**
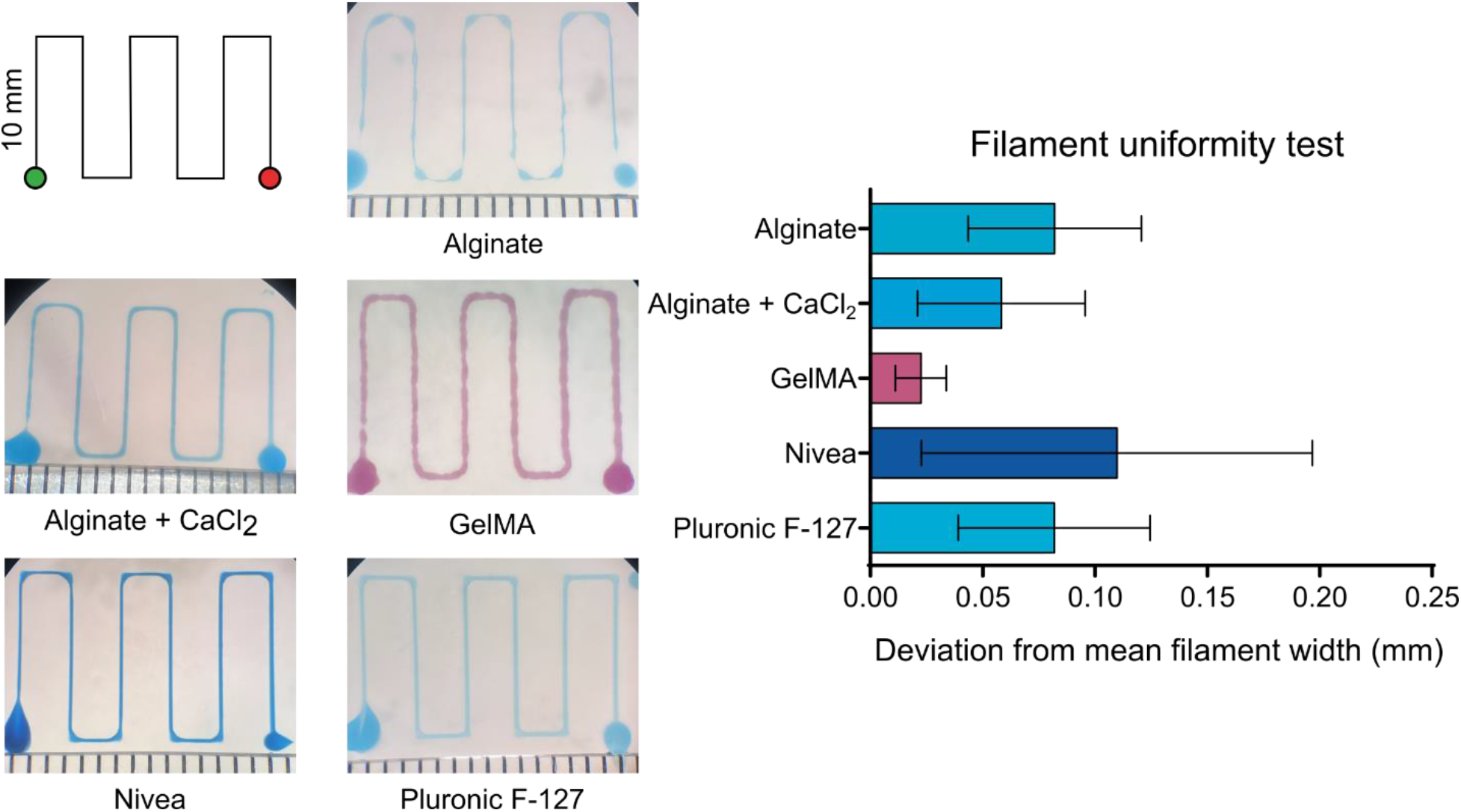
A filament uniformity test reveals that filament uniformity is consistent across different biomaterial inks. For this figure and following figures, green indicates the start position of the needle and red indicates the end position of the needle.

**Figure 23:**
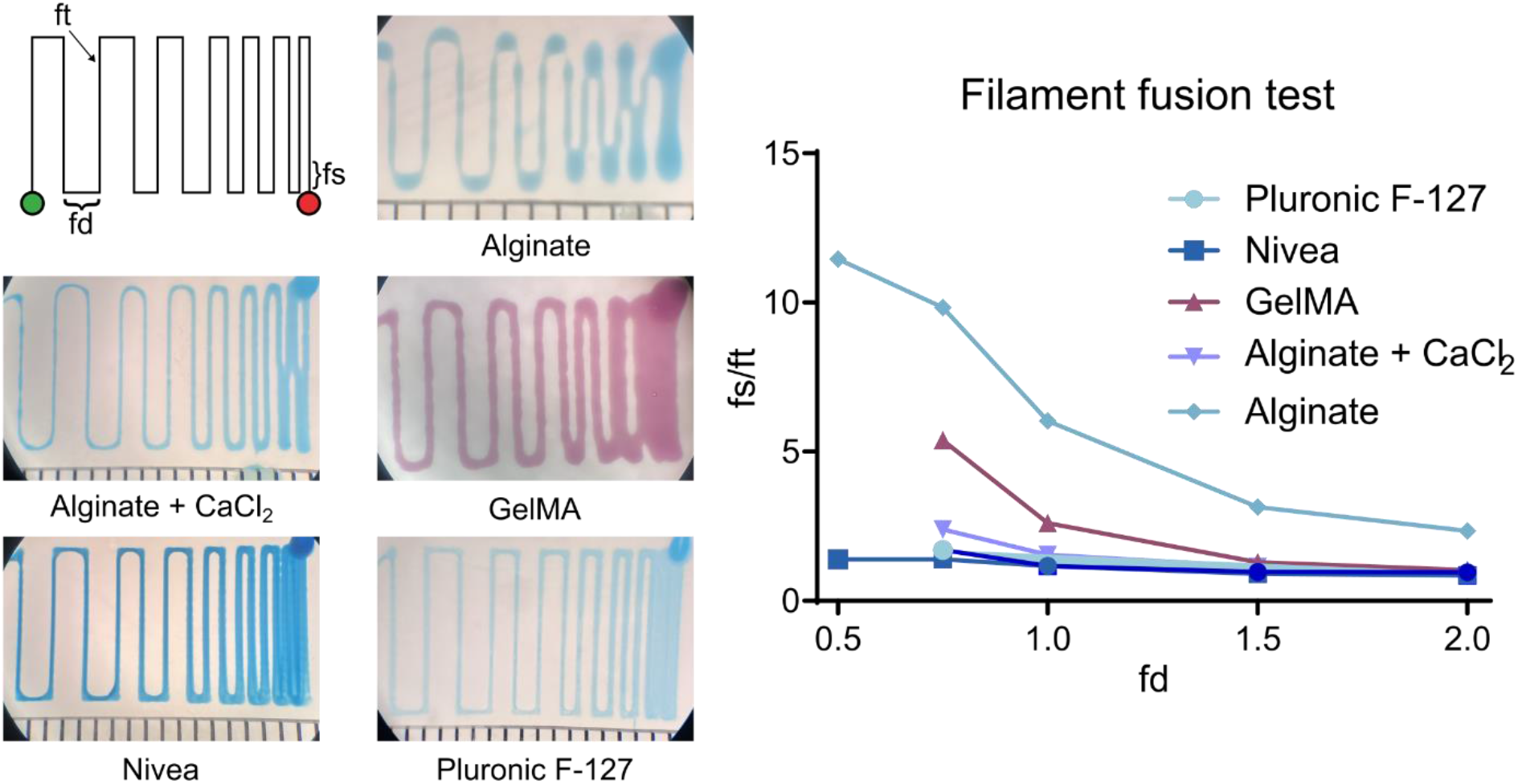
A filament fusion test revealed that most inks are not susceptible to a loss of resolution due to the surface tension of a nearby strand of filament. When the distance between strands (fd) was less than 1 mm, both GelMA and fluid alginate tended to merge into a single filament (as measured by the fused filament length (fs), normalized by the filament thickness (ft)). Missing data points indicate that adjacent lines merged at this filament spacing distance and thus a fs value could not be calculated.

**Figure 24:**
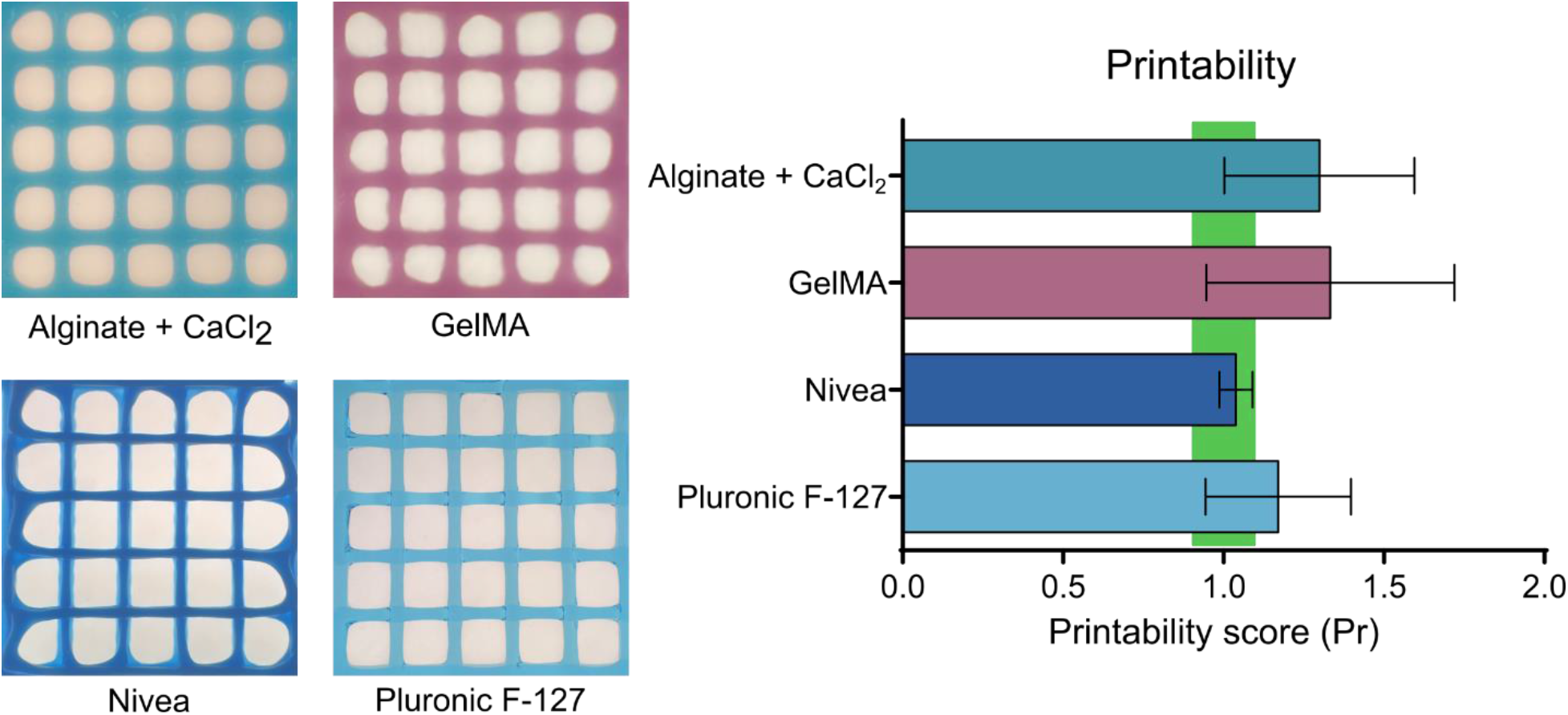
To assess printability, filaments were printed on the Enderstruder in a 5 x 5 grid, with each pore consisting of a 1 mm x 1 mm square. All exhibited printability as previously defined by Ouyang *et al.* (green box, 0.9 < P < 1.1).

**Figure 25:**
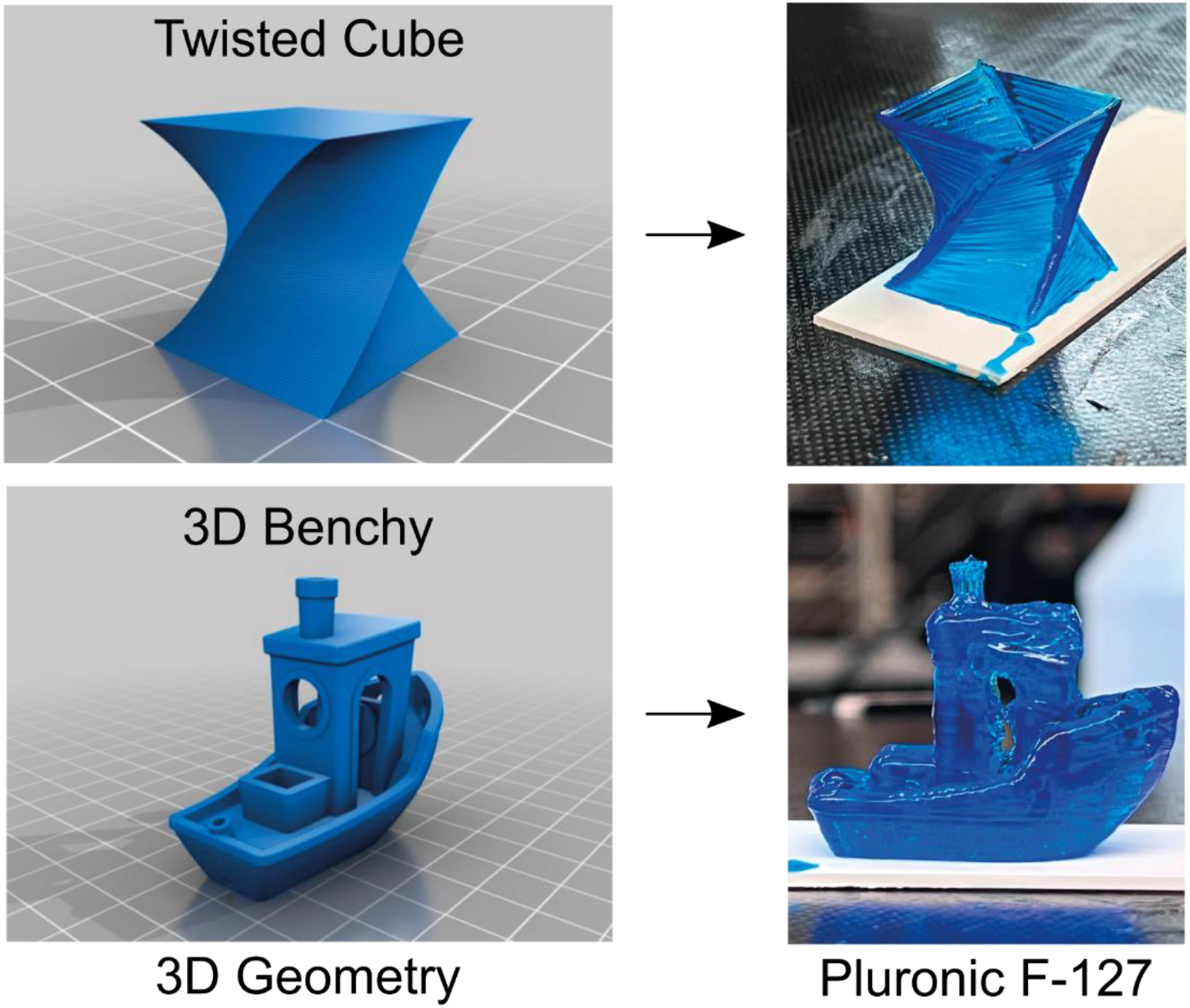
The Enderstruder was able to print the geometries of a twisted cube and a 3D “Benchy” at high resolution from Pluronic F-127.

## CRediT author statement

Domenic J. Cordova: Conceptualization, Methodology, Software, Validation, Investigation, Writing – Review & Editing. Angel A. Rodriguez: Conceptualization, Methodology, Software, Investigation, Original Draft Preparation, Visualization, Writing - Review & Editing. Sabrina C. Woodward: Conceptualization, Software, Validation, Formal Analysis, Writing - Review & Editing. Cody O. Crosby: Conceptualization, Resources, Original Draft Preparation, Visualization, Supervision, Project administration, Funding acquisition, Writing - Review & Editing

## Acknowledgments

The authors would like to thank Kristie Cheng, B.S., Noor Nazeer, and Gabriela Nicole Hislop Gomez of Southwestern University for synthesizing and loading the biomaterial inks and FRESH support baths used in the Validation section of this article. The authors would also like to acknowledge the contributions of Katie Rosenkrantz, B.S., and Andrew Vergote, B.S., for their prior work on E-step calibration and developing the suspension bath protocol for the lab, respectively. The corresponding author would like to thank Dr. Nancy Chick, Dr. Myra Monreal, Angie Dewberry, M.A., and Dr. Amber Reed for their feedback while drafting the manuscript through an Associated Colleges of the South (ACS) writing accountability group (WAG). Figure 1 and Figure 5 were made with photographs taken by Todd White, Southwestern University’s videographer. All figures were edited with the aid of Estefania Pedrazas.

Funding: This work was supported by the Robert A. Welch Foundation (AF-0005), the Sam Taylor Foundation, and a generous gift to Southwestern University from Bob and Annie Graham.

During the preparation of this work the authors used ChatGPT to improve the flow and clarity of the manuscript. After using this tool, the authors reviewed and edited the content as needed and take full responsibility for the content of the publication.

